# Modeling the age distribution of retinoblastomas

**DOI:** 10.1101/2021.11.24.469838

**Authors:** Alexandr N. Tetearing

**Affiliations:** St. Petersburg State University

## Abstract

In this work, based on real data on the size of the eyeball (in a fetus, in a child, and in young people under 20), we constructed a model function of the growth of the retinal cell tissue.

We used this function to construct a theoretical age distribution of retinoblastomas. We constructed theoretical age distributions for four different models of retinoblastoma: a complex mutational model, a third mutational model, a model with a sequence of key events, and a model of a single oncogenic event with two different latencies (hereditary and non-hereditary retinoblastoma).

We compared the theoretical age distribution of retinoblastomas with the real age distribution based on SEER data (Surveillance Epidemiology and End Results; program of the American National Cancer Institute). In total, we examined 843 cases in women and 908 cases in men.

For all models (separately for women and men), we obtained estimates of the following cancer parameters: the specific frequency of key events (events that trigger cancer); the duration of the latency period of cancer; the number of key events required for cancer to occur.

For the composite age distributions, we calculated the theoretical mean age at diagnosis for hereditary and non-hereditary retinoblastomas.

The best approximation accuracy (for male and female forms of retinoblastoma) is shown by a model with a sequence of key events.

## 1. Introduction

Retinoblastoma (childhood malignant tumour of the retina) differs from other cancers in that the peak incidence occurs in the first years of a child’s life.

About half of retinoblastoma cases are hereditary – patients have a parent with this disease. In [1], Alfred Knudson suggested that retinoblastoma occurs when both alleles of the retinoblastoma gene are damaged as a result of mutation (the tumour suppressor gene Rb was discovered later, in 1986 [2]). In hereditary retinoblastoma, the patient receives one mutant allele from the parent, the second, later mutation, occurs by chance. In the non-hereditary form of the disease, both alleles are damaged by random mutations.

Despite the long period of study of retinoblastoma, the picture of the onset of the disease is not completely clear. It is known that not all patients with non-hereditary retinoblastoma have mutations in the Rb gene [3].

In this work, we will try to obtain a theoretical form of the age distribution of retinoblastomas for different models of retinoblastoma occurrence (different models give different age distribution) and compare them with real age distributions.

## 2. Cellular tissue growth function

### 2.1. Initial data

To construct the age distribution of the incidence of retinoblastoma, we, as a rule, need a certain model function of the growth of cell tissue.

As the initial data for constructing the function of cell tissue growth (retinal cell tissue), we used data on the diametral size of the eyeball.

The size of the eyeball is measured both in utero (in the fetus) and in the postpartum period (in a child) by computed NMR tomography (nuclear magnetic resonance), ultrasound scanning (US) and infrared interferometer. Research is carried out to monitor various eye diseases (myopia, glaucoma, cataracts). Usually, the transverse diameter of the eye and the axial length (the distance between the anterior and posterior walls of the eyeball) are measured.

To construct the retinal growth function, we used data obtained from three different sources: [4],[5],[6].

In [4], measurements of the diameter of the fetal eye at the age from 11 weeks of gestation to childbirth, performed by NMR tomography, are presented: 127 cases are presented with known age in weeks. For each case, the average diameter of the left and right eye is presented.

The work [5] is a review article summarizing the work devoted to the measurement of the axial length of the eye in children from 0 to 3 years old, performed by the method of ultrasound scanning. The authors analyzed the measurements of 6575 eyes presented in 27 scientific articles. The data are categorized by age group and the mean is presented for each group. The data were not separated by sex of the child (this is not essential in infancy).

The work [6] presents the results of measurements of healthy eyes (238 cases), performed using an infrared laser interferometer in patients from 4 to 20 years old (separately for men and women). Widely scattered data, divided into 7 age groups (the average value for the group is given).

The three datasets differ in the measurement method. In the fetus, before birth, measurements are carried out on an NMR tomograph. This is bulky and expensive equipment (it takes a small room). Therefore, less data is collected than data from the other two age groups. From birth to 4 years of age, eye size is measured using an ultrasound scan. This is a desktop equipment, quite cheap and widely used, and this group of measurements is represented by the largest amount of data. The third group of measurements – children from 4 years old – is represented by data obtained on a laser scanning interferometer. This is a tabletop device that measures the axial length of the eyeball with a low-power laser beam through the open eye pupil. The child should consciously look into the eyepiece of the device for several seconds. For this reason, measurements are taken on children aged 4 years and older. The interferometer provides greater measurement accuracy compared to ultrasound, but measurements in young children are not made on it.

In order to construct a function of growth of cellular tissue (retina) on the basis of these data, we must calculate the volume of cellular tissue and the number of cells in a given volume from the linear size of the eye (eye diameter *D*).

Assuming that the retinal cells are located on the inner spherical surface of the eye in one layer, we can also assume that the total number of cells *N*(*t*) (for an adult organ and for an organ in the process of growth) is proportional to the area of a sphere with diameter *D*:

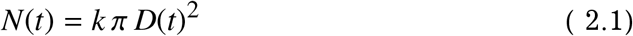

Here *k* is some constant coefficient that can be set if we know the number of cells in the tissue of an adult organ. Eye diameter *D*(*t*) changes with time during growth.

At present, it is not known exactly what type of retinal cells the tumour develops from. It is assumed [7] that the tumour originates from the progenitor cells of the retinal light-sensitive cones.

In [8], the number of cone cells in the retina of an adult eye is estimated at 4.6 million cells. Therefore, we assume that the number of cells in the cellular tissue is 9.2 · 10^6^. We set the coefficient *k* in (2.1) so that the value of the function *N*(*t*) in adulthood is equal to 9.2 · 10^6^ cells.

The set of discrete points *N_i_* of the cell tissue growth function obtained by formula (2.1) based on real data sets on the linear dimensions of the eyeball is shown in Fig. (2.1). The figure shows data from all three sources: [4],[5],[6]. Data from work [5] (children from 0 to 4 years old) are marked with black squares. The top graph shows the data [4], [5] for ages from 3 to 33 months (data without gender division). The middle graph (different timescale) shows data for ages from 0 to 240 months (20 years) for women. The bottom graph shows data for men aged 0 to 240 months (20 years). Time is counted from the beginning of gestation. The time of birth *t*_0_ is marked with a vertical dashed line in all three graphs.

**Figure 2.1.**
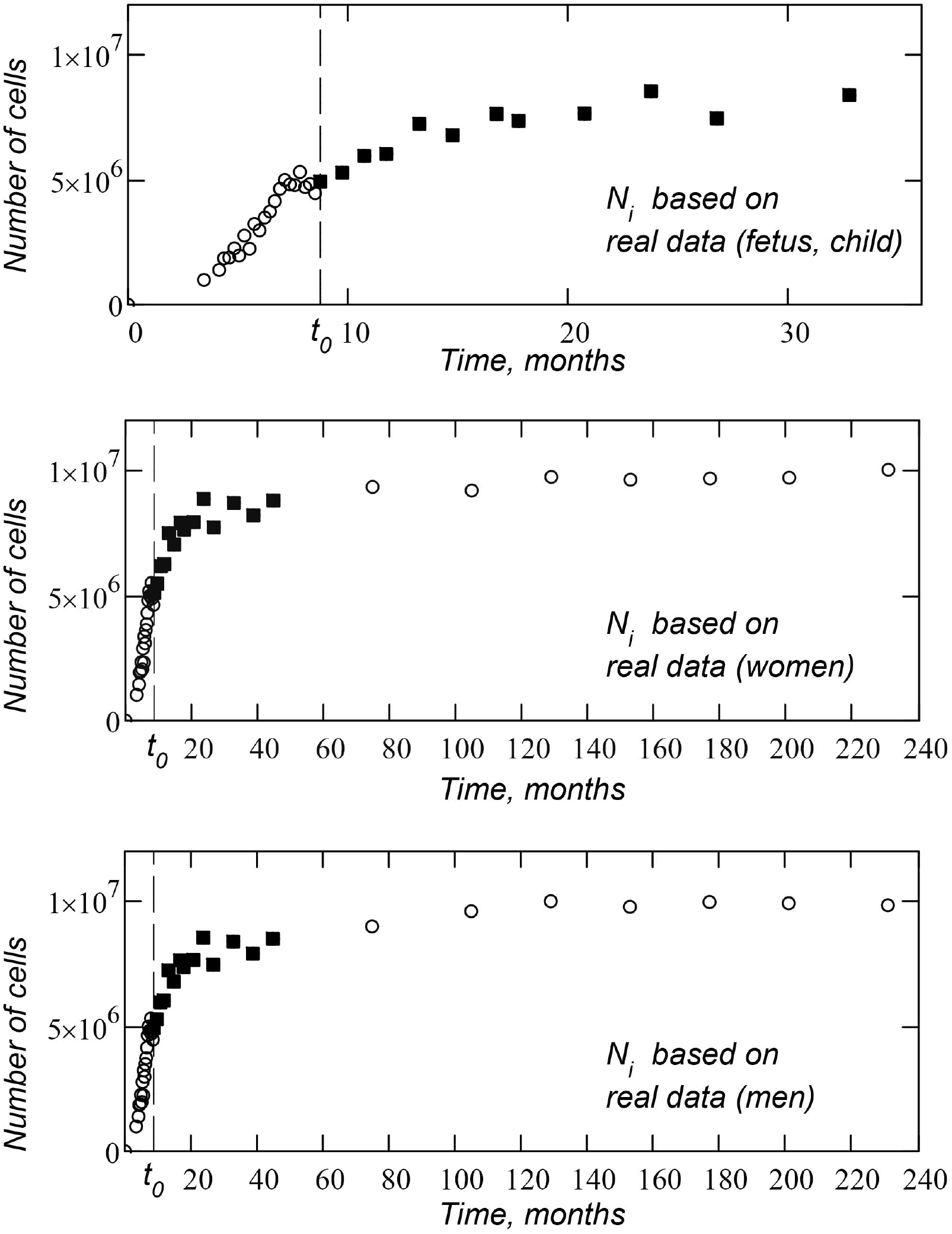
Retinal cell growth (plot points are based on real data). The upper figure is data for ages from 3 to 33 months. Average figure is data for ages from 3 to 240 months (20 years) for women. Bottom figure is data for ages from 3 to 240 months (20 years) for men. Time is counted from the start of gestation. The vertical dashed line is the time of birth *t*_0_.

We will approximate this data using a theoretical growth function (presented in the next section). After performing the first approximation, we can establish the exact value of the coefficient *k* in formula (2.1). In Fig. 2.1 the coefficient *k* is *k* = 5.87 · 10^3^ for women and *k* = 5.66 · 10^3^ for men.

Besides the real dataset, we set the initial point of the growth function *N*(0) = 1. Here we assume that at the initial moment of time *t* = 0 the cellular tissue consists of one cell.

### 2.2. Model function of growth

As a growth function, we use our usual growth function, which we used earlier – for example, in [9]:

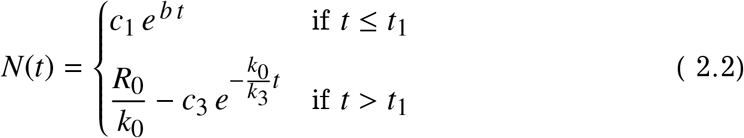

This is the simplest growth function, and, in this case, *c*_1_ = 1 (at the initial moment of time *t* = 0, the tissue consists of one cell).

The function consists of two pieces (of two curved lines) given at intervals [0; *t*_1_] and [*t*_1_; ∞]. On the first interval, the function *N*(*t*) grows exponentially, after time *t* = *t*_1_, the function *N*(*t*) encounters some growth-limiting factors, and the growth of the function slows down, exponentially approaching a certain limit:

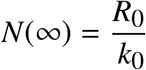

In our case:

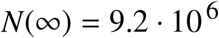

The *R*_0_/*k*_0_ ratio specifies the number of cells in an adult organ. The constant *b* from the upper equation of system (2.2) is determined by the expression:

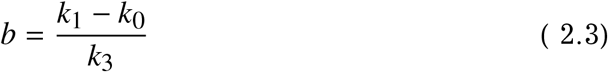

The parameters *t*_1_, *R*_0_ and *k*_3_ are related by the equation:

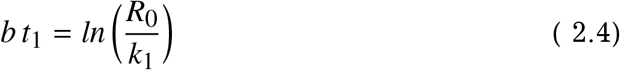

The constant *c*_3_ (this is the constant of integration) is found from the equation:

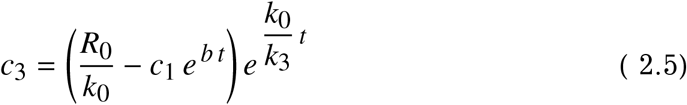

All parameters included in equations (2.2) are described in detail in the book [10].

### 2.3. Approximation of the initial data

In this section, based on real data sets, we will construct *N*(*t*) function (model function of cell tissue growth), which we will use later to obtain the age distribution of retinoblastomas.

We need to substitute into the equations describing this model function (2.2) such (optimal) coefficients *R*_0_, *k*_0_, *k*_3_ in order to minimize the value of the error function *Er*

We need to substitute certain coefficients *R*_0_, *k*_0_, *k*_3_ in the equations describing this model function (2.2) to minimize the error function *Er* (least square method):

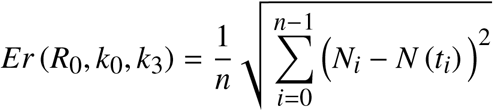

Optimization of the error function is carried out programmatically, using the quasi-Newtonian method.

Figures 2.2 and 2.3 show graphs of cell tissue growth functions obtained as a result of approximation (separately for women and men). The growth function *N*(*t*) is shown by the bold solid line. The datasets are shown with open circles.

**Figure 2.2.**
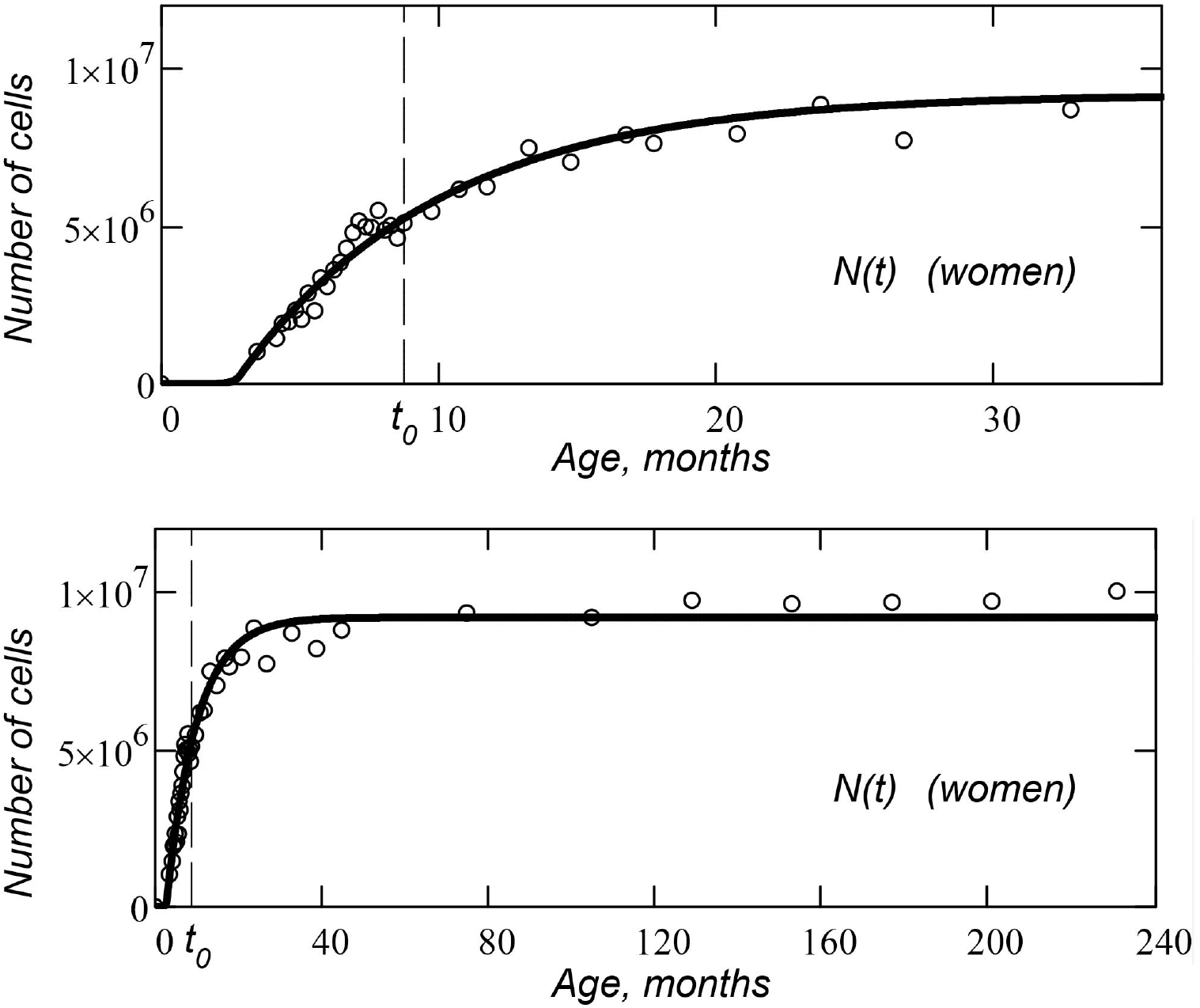
Retinal cell growth function (bold solid line) is approximation for a real dataset (women). Real data are shown with open circles. The top and bottom figures show the graph at different time scales. Age is counted from the beginning of gestation. The vertical dashed line shows the time of birth *t*_0_.

**Figure 2.3.**
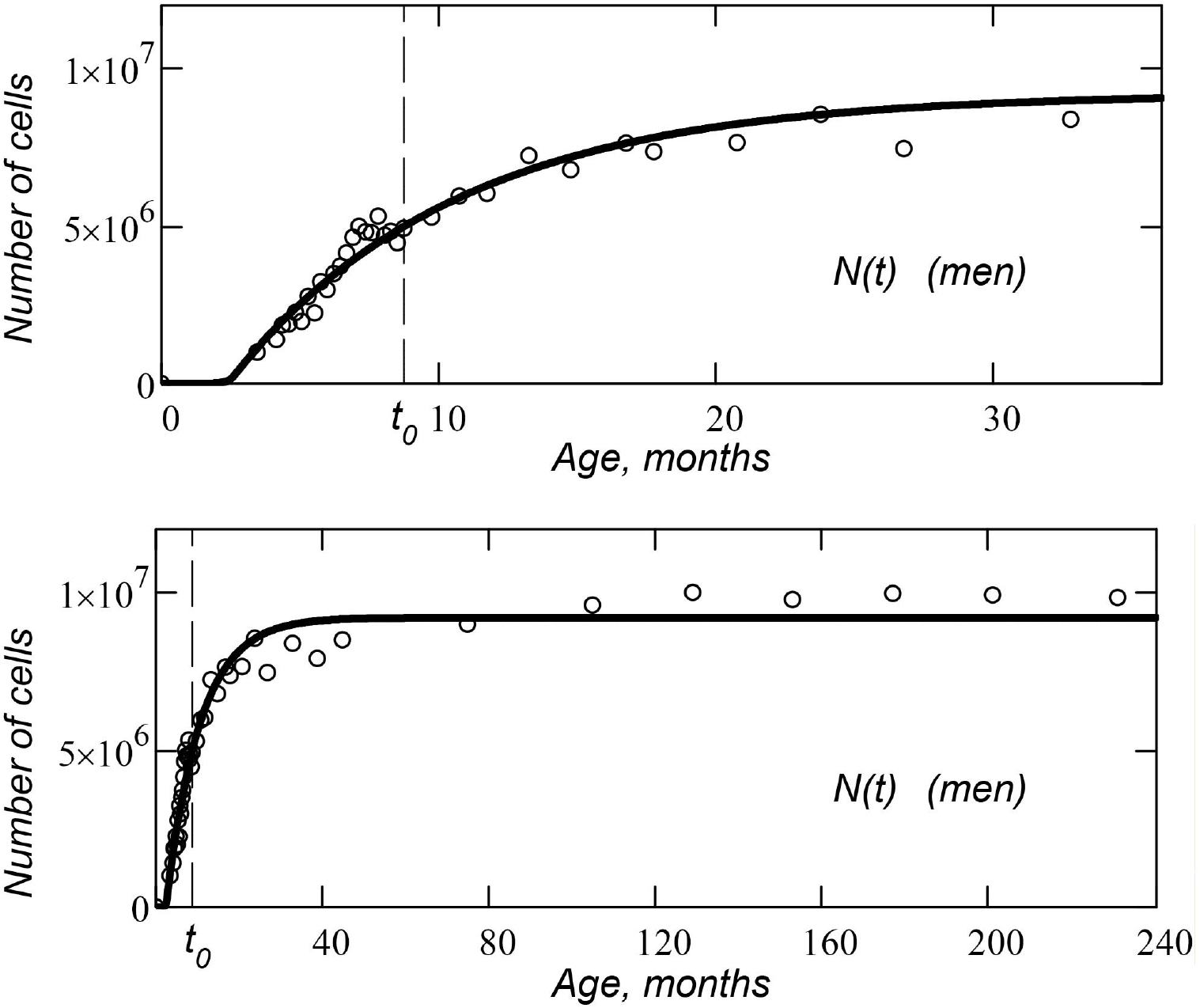
Retinal cell growth function (bold solid line) is approximation for a real dataset (men). Real data are shown with open circles. The top and bottom figures show the graph at different time scales. Age is counted from the beginning of gestation. The vertical dashed line shows the time of birth *t*_0_.

The top and bottom plots in Figures 2.2 and 2.3 show the same function at different time scales. Age is measured from the start of gestation. The vertical dashed line shows the time of birth *t*_0_.

As a result of optimization, we obtained the following values of the parameters of the cell tissue growth function: *R*_0_ = 2.78 · 10^5^, *k*_0_ = 0.30, *k*_3_ = 0.22, *k* = 5.87 · 10^5^, *t*_1_ = 2.8 (women); *R*_0_ = 2.32 · 10^5^, *k*_0_ = 0.25, *k*_3_ = 0.20, *t*_1_ = 2.6 (men). The coefficient *k*_1_ in both cases is considered equal to one. The root-mean-square error of approximation is *Er* = 7.04 · 10^4^ for women and *Er* = 7.99 · 10^4^ for men.

We will use these two functions (with the obtained optimal coefficients) to construct the age distribution of retinoblastomas.

## 3. Real age distribution of retinoblastomas. Data preparation

To construct the real age distribution of retinoblastomas, we used data from the SEER program (Surveillance Epidemiology and End Results) of National Cancer Institute (US) for the period from 2000 to 2016 [11]. The SEER database contains information on 843 cases of retinoblastoma in women and on 908 cases in men.

The preparation of cancer incidence data is described in detail in [12] and [13], so here we only give a general description of the algorithm.

First, from the database, we select all cases of diseases (separately for women and men) for one year – let’s say we want to select cases of female retinoblastoma in 2000. We sort all cases for 2000 by age.

We divide the number of cases for each age by the US population for that age and gender (US Census Bureau data [14]). Next, we divide this result by 0.28 because SEER data covers 28 percent of the US population. [15]. The resulting number is the incidence of retinoblastoma per one person of a given age and gender. For each age (age is taken in 1 year increments) we get a certain number.

We carry out similar actions for each of the next 16 years (each year separately).

For each age, we summarize 17 incidence rates obtained for each year (from 2000 to 2016). We divide the sum by 17 and multiply by 100,000. Thus, we get the incidence for a given age per 100,000 people (separately for women and men). The numbers (incidence cases) obtained for each age are plotted on the general graph of the age distribution of incidence.

The resulting graphs of the real age distribution of retinoblastomas per 100,000 people (separately for women and men) are shown in Fig. 3.1.

**Figure 3.1.**
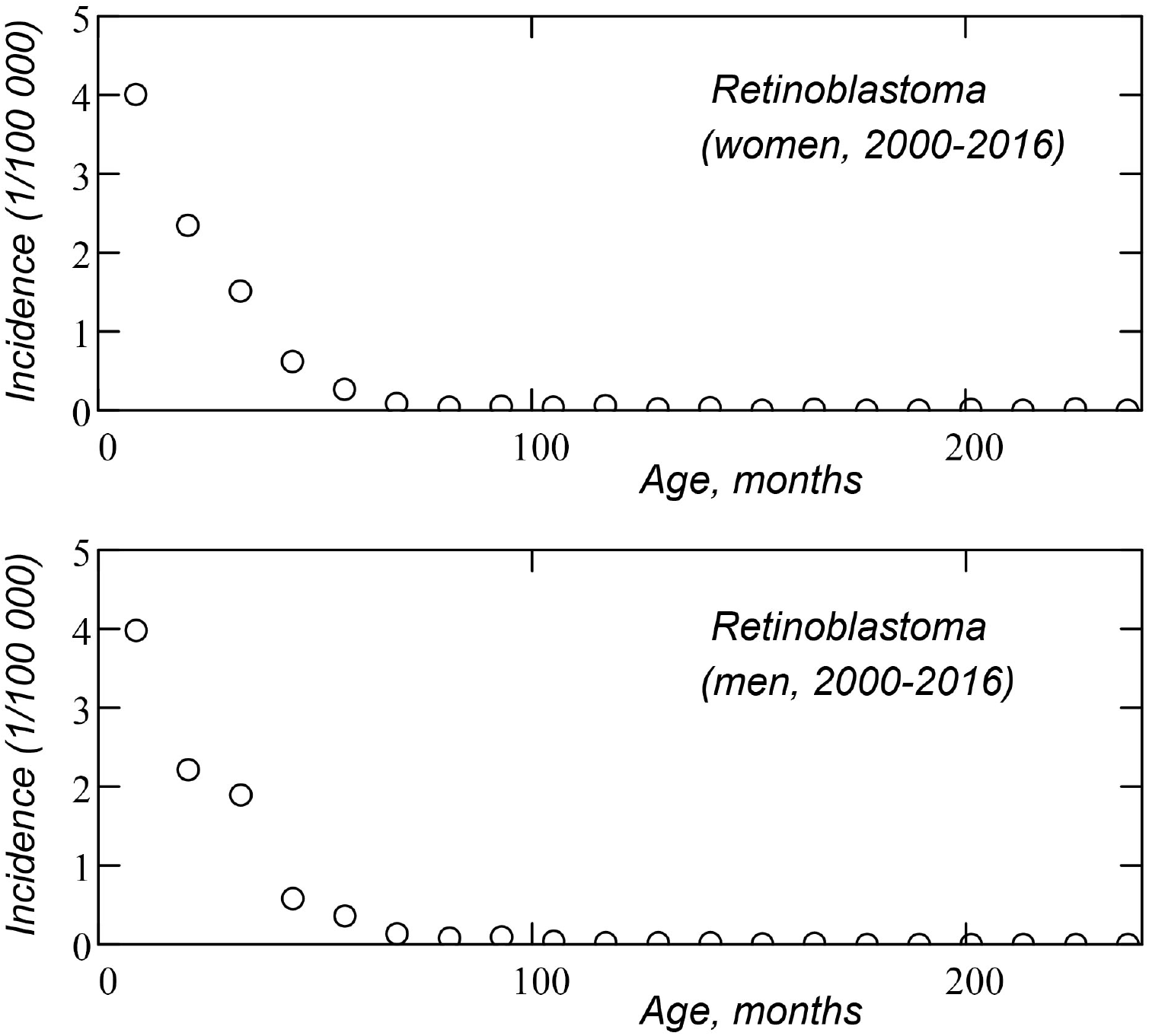
Age distribution of retinoblastomas (real data) per 100,000 people for women (upper graph) and men (bottom graph) for the period from 2000 to 2016.

## 4. Complex mutational model

### 4.1. Age distribution functions

As a model of oncological disease, we use the mutational model presented in [9]. According to the model’s definition, cancer occurs when a cell receives a certain number of mutations (key events). We will consider age distributions for the case of one mutation per cell, two mutations per cell, as well as a composite distribution combining (in some ratio) cases with one mutation and cases with two mutations per cell.

The theoretical equations for the age distribution functions are as follows:

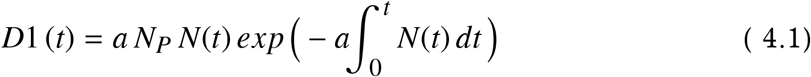

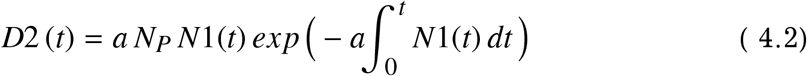

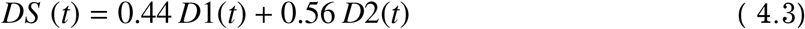

The function *D*1(*t*) is the age distribution of retinoblastomas, built for the case when the disease occurs after one mutation in the cell; *D*2(*t*) function is the age distribution constructed for the case when the disease occurs after two mutations in the cell; *DS*(*t*) function is the composite age distribution obtained for the composite function *NS*(*t*) (see below).

In these equations, *a* is the average frequency of specific mutations (the average number of mutations occurring in a cell per unit of time). The constant *N_P_* is the number of people in the group for which the age distribution is constructed. *N*(*t*) function is the cell tissue growth function (the function described in the previous section). The *N*1(*t*) function describes the number of cells (in the considered anatomical tissue) that have one or more mutations.

*DS*(*t*) function reflects the case when the age distribution of retinoblastomas is formed by two different groups of patients. The first group gets cancer after one cell mutation (inherited form of retinoblastoma), the second group gets the disease after two mutations in the cell (non-inherited form of retinoblastoma). The proportion of patients in the first group is 44 percent, the proportion of patients in the second group is 56 percent of the total size of the group, for which the age distribution of retinoblastomas is constructed (this is 44 and 56 percent, respectively, of *N_P_* constant).

We chose the percentage ratio between the two groups with different numbers of mutations initiating the disease based on the data presented in [16], where 918 cases of retinoblastoma were analyzed. According to the results of the study, the proportion of diseases with documented hereditary form of retinoblastoma was 44 percent. The authors note that the proportion of hereditary diseases was significantly higher than is usually indicated in other scientific articles. Therefore, when identifying this ratio, it is important to look at the total number of cases considered.

The function *N*1(*t*) (the case when a tumour is formed from two mutations in a cell) is given by the equation:

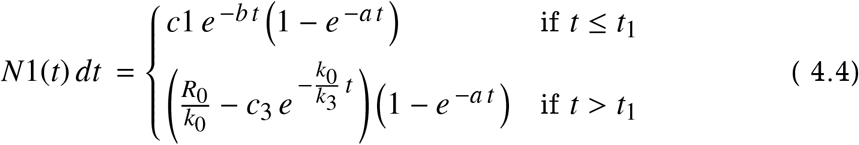

The functions *N*(*t*), *N*1(*t*) and *NS* (*t*) are piecewise functions that are given by different equations on two different time intervals (from 0 to *t*_1_ and from *t*_1_ to infinity). Therefore, after the time *t*_1_, the integration in formulas (4.12-14) should be carried out separately for each time interval. For example, for *N*1(*t*) function it will look like this:

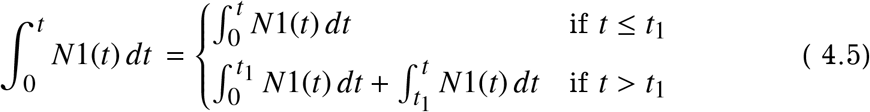

*N*1(*t*) function on the interval [0; *t*_1_] and the *N*1(*t*) function on the interval [*t*_1_; ∞] are two different functions. Therefore, non-observance of this rule will lead to a discontinuity of the function *D*1(*t*) at the point *t* = *t*_1_.

### 4.2. Approximation method

We perform an approximation of the real age distribution (Figure 3.1) using theoretical functions (4.1), (4.2), (4.3).

In the real age distribution, cases of diseases are grouped into age groups with a wide range of ages (the width of the group is one year or 12 months). The theoretical age distribution function varies greatly within one year (we usually consider the graph of the theoretical function in months, not years). Therefore, for comparison with the real age distribution, we must present your theoretical distribution in the form of a histogram (bar chart), where each bar represents the total number of cases that fell into a given age group from our age distribution. The age groups on our histogram will be wide – the width is one year or more (for the zero group) – this corresponds to the data of the real age distribution of retinoblastomas.

The height of the histogram bar is the integral of the theoretical age distribution function. For example, for the zero bar and the age distribution function *D*1(*t*) we have:

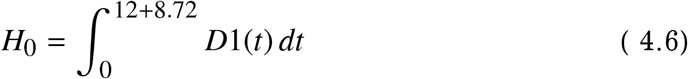

The height of the zero column of the histogram is the number of cases of retinoblastoma that occurred in the considered group of people in the age interval from 0 to 12 months. In this case, the time *t* (age) is counted from the beginning of pregnancy. We assume that the pregnancy lasts 266 days (38 weeks) from the time of fertilization of the mother’s egg, and therefore the time of birth on the scale of months is *t* = 8.72. An infant 11 months old falls into the *H*_0_ group of the histogram (since its age is 0 completed years from birth).

The height of the first bar of the histogram is:

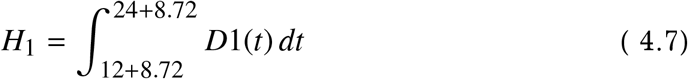

This age group includes infants between the ages of 12 and 24 months (babies who are 1 full year old).

The height of the second bar of the histogram is:

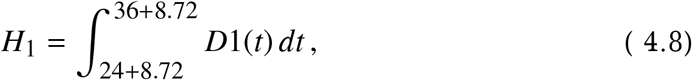

etc.

The height of the last bar of the histogram is:

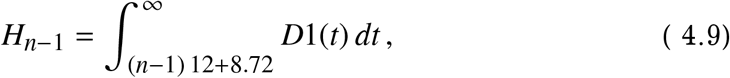

In our case, this column can be considered equal to zero, since retinoblastoma is a disease of an early age, and the number of cases of retinoblastoma registered after 80 years is zero.

The age distribution of *D*2(*t*) (the case when the disease arises from two mutations in the cell) and the histogram of this distribution are shown for example in Fig. 4.1. Note that the graphs have different scales on the vertical axis.

**Figure 4.1.**
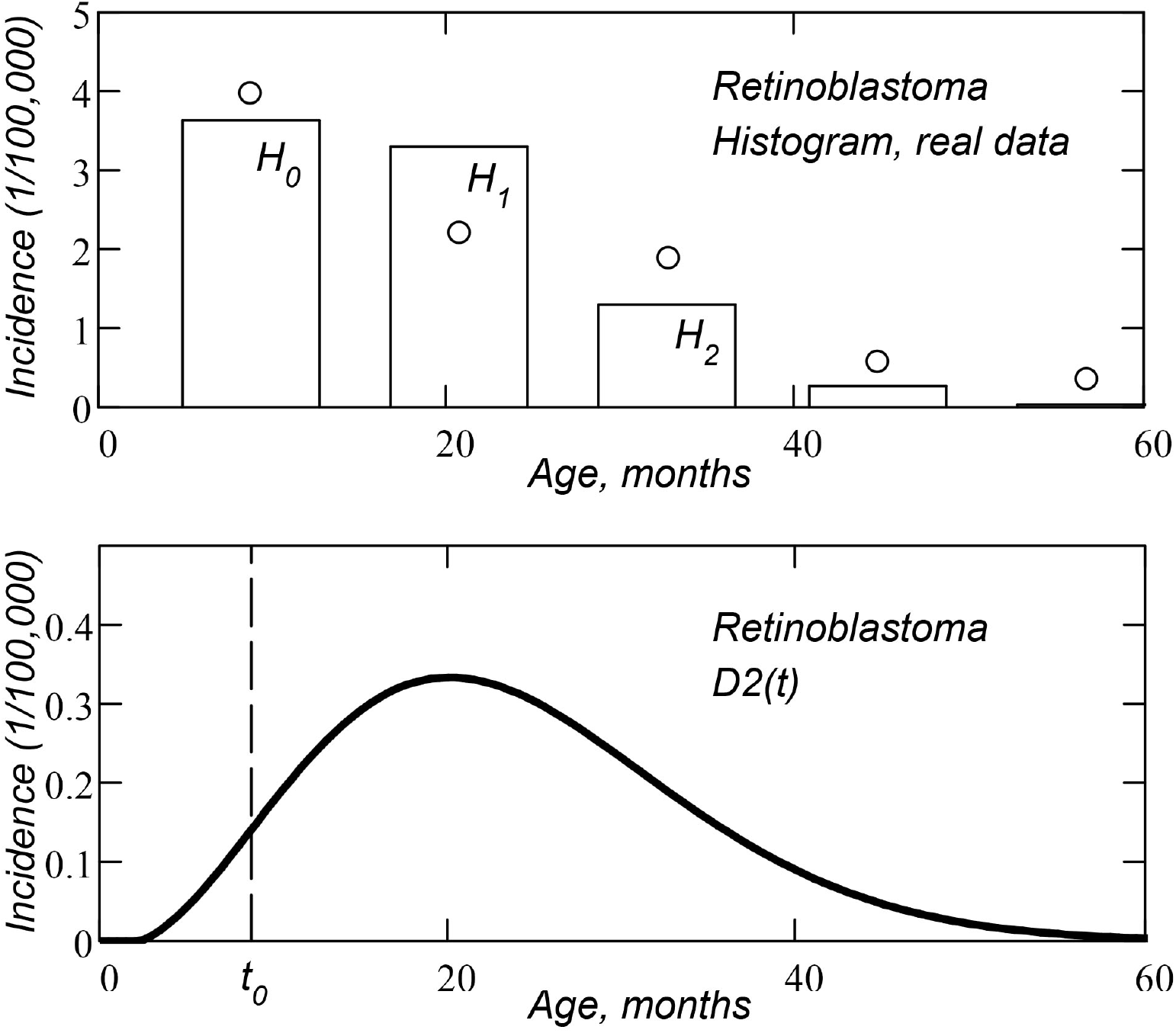
Histogram *H_i_* of age distribution of retinoblastomas (rectangular bars, top graph) and real age distribution data (open circles, upper graph); age distribution function *D*2(*t*) (bold curve line, bottom graph) for the case when the disease occurs due to two mutations (key events) in the cell.

When constructing the graphs, we used the cell tissue growth function *N*(*t*) (discussed above and built on the basis of real data on the size of the eyeball in men) and the following parameters: *a* = 2.0.^-5^, *N_P_* = 8.5. *T_S_* = 0, *z* = 2.

The histogram of the age distribution of the incidence obtained by the above method is used as a discrete approximation function to approximate the real data set of the age distribution of retinoblastomas.

Choosing the optimal parameters *a, N_P_, T_S_* (parameters of the age distribution function used to construct the histogram), we minimize the error function *Er* (least squares method):

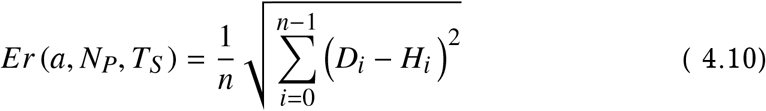

Here *D_i_* is a value from the real age distribution; *H_i_* is the value of the *i*-th bar of the histogram built for the theoretical age distribution; *n* is the number of points in the real dataset.

Together with the parameters *a* (the average number of mutations occurring in the cell per unit of time) and *N_P_* (the number of people in the considered group, for which the age distribution is constructed), we also optimize the parameter *T_S_* – this is the time shift between the onset of the disease (the key event initiating the disease) and the detection of cancer (the latency period of the disease). For example, if *T_S_* = 2, we shift the theoretical age distribution function 2 months to the right in the timeline.

We carry out the approximation of the data set programmatically, using the quasi-Newtonian method.

In addition to the absolute error function *Er* for the found optimal approximation parameters, we calculate the relative error function *Er*%:

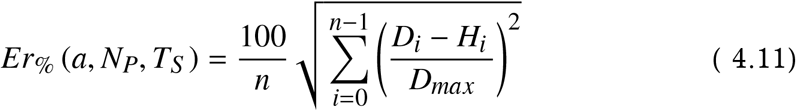

The function shows the relative root mean square error of approximation as a percentage of the maximum value of *D_max_* from a set of real data.

### 4.3. Calculating the mean age at diagnosis

For the compound functions of the age distribution of morbidity (when the group in question includes patients with hereditary and non-hereditary forms of retinoblastoma), it would be interesting to determine the average age at the time of diagnosis in order to compare the theoretical age with the real one.

If we know the age distribution function *DS*(*t*), then the average time (age) of the diagnosis can be found as the projection onto the time axis of the point that is the center of mass of the figure bounded by *DS*(*t*) function and time axis. Here we assume that the mass of the figure is proportional to its area.

It is also necessary to take into account the fact that all cases of a disease that began before birth can be diagnosed only after birth (not earlier). Therefore, we calculate the average age of the diagnosis *t_d_* in two stages. First, we find the projection of the center of mass of the figure, including all cases of the disease for ages from *t*_0_ (time of birth) to infinity. This age *t*_*d*0_ is found from the equation:

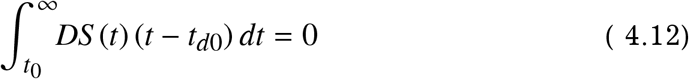

Since we calculate age from the time of fertilization of the mother’s egg, the time of birth *t*_0_ is 8.72 months.

Then we find the center of mass *t_d_* of two material points with different masses: the first point is located on the time axis at the point *t* = *t*_0_ and has a mass equal to the sum of all cases of the disease that occur before birth, the second material point has a mass equal to the sum of all cases that occur after birth. This mass is located on the time axis at the point *t* = *t*_*d*0_.

Thus, the point *t_d_* (average age at diagnosis) is found from the equation:

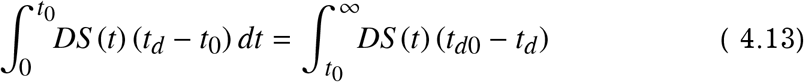

It should be remembered that the mean age at diagnosis is not the time of the maximum value of the age distribution (distribution peak time – the age for which the maximum number of cases of the disease is recorded). Also, the mean age at diagnosis is not the median age of the age distribution. (age that divides the number of cases into two equal halves – the number of cases reported before the median age is equal to the number of cases reported after the median age).

### 4.4. Results

For the mutational model of retinoblastoma (separately for men and women), we performed software optimization of the model parameters for cases when the disease develops as a result of one mutation (key event) in the cell; for the case when the disease develops as a result of two mutation in the cell; and for the composite case where 44 percent of patients (in the group for which the age distribution is constructed) develop the disease as a result of one mutation in the cell (hereditary retinoblastoma), and 56 percent of patients develop the disease as a result of two mutations in the cell (non-hereditary retinoblastoma).

As already mentioned, the specified percentage was selected based on the data from [16].

The optimization results are presented in Table 4.1 for three cases with different numbers of initiating mutations. The *z* parameter is the number of mutations (key events) in the cell that initiates the disease. The parameter *z* = 1.56 refers to the composite age distribution (44 percent of the group are patients with one mutation, 56 percent of the group are patients with two mutations.).

**Table 4.1.** Optimal approximation parameters for mutational model of retinoblastoma.

| **Retinoblastoma** | $T_S$ | $a$ | $N_P$ | $Er$ | $Er\%$ |
| --- | --- | --- | --- | --- | --- |
| Female; $z=1$ | 1.6 | $6.3 \cdot 10^{-9}$ | 9.1 | $3.41 \cdot 10^{-3}$ | 0.09 |
| Female; $z=1.56$ | 10.9 | $2.2 \cdot 10^{-5}$ | 8.4 | $4.65 \cdot 10^{-3}$ | 0.12 |
| Female; $z=2$ | 0 | $2.1 \cdot 10^{-5}$ | 7.8 | $12.58 \cdot 10^{-3}$ | 0.32 |
| Male; $z=1$ | 0.4 | $5.4 \cdot 10^{-9}$ | 9.6 | $7.18 \cdot 10^{-3}$ | 0.18 |
| Male; $z=1.56$ | 12.2 | $2.2 \cdot 10^{-5}$ | 8.7 | $4.58 \cdot 10^{-3}$ | 0.12 |
| Male; $z=2$ | 0 | $2.0 \cdot 10^{-5}$ | 8.0 | $16.75 \cdot 10^{-3}$ | 0.42 |

The *T_S_* parameter is the latent period of the disease (the time from the key event that initiates the disease to the diagnosis).

The a parameter is the specific frequency of key events – the average number of key events that occur in a group of cells per unit of time (month).

The parameters *Er* and *Er*_%_ are absolute and relative mean square error. The relative error is calculated as a percentage of the maximum value from the set of real data on the age distribution of retinoblastomas.

The obtained functions with optimal approximation parameters are shown in Figures 4.2, 4.3, 4.4, and 4.5.

**Figure 4.2.**
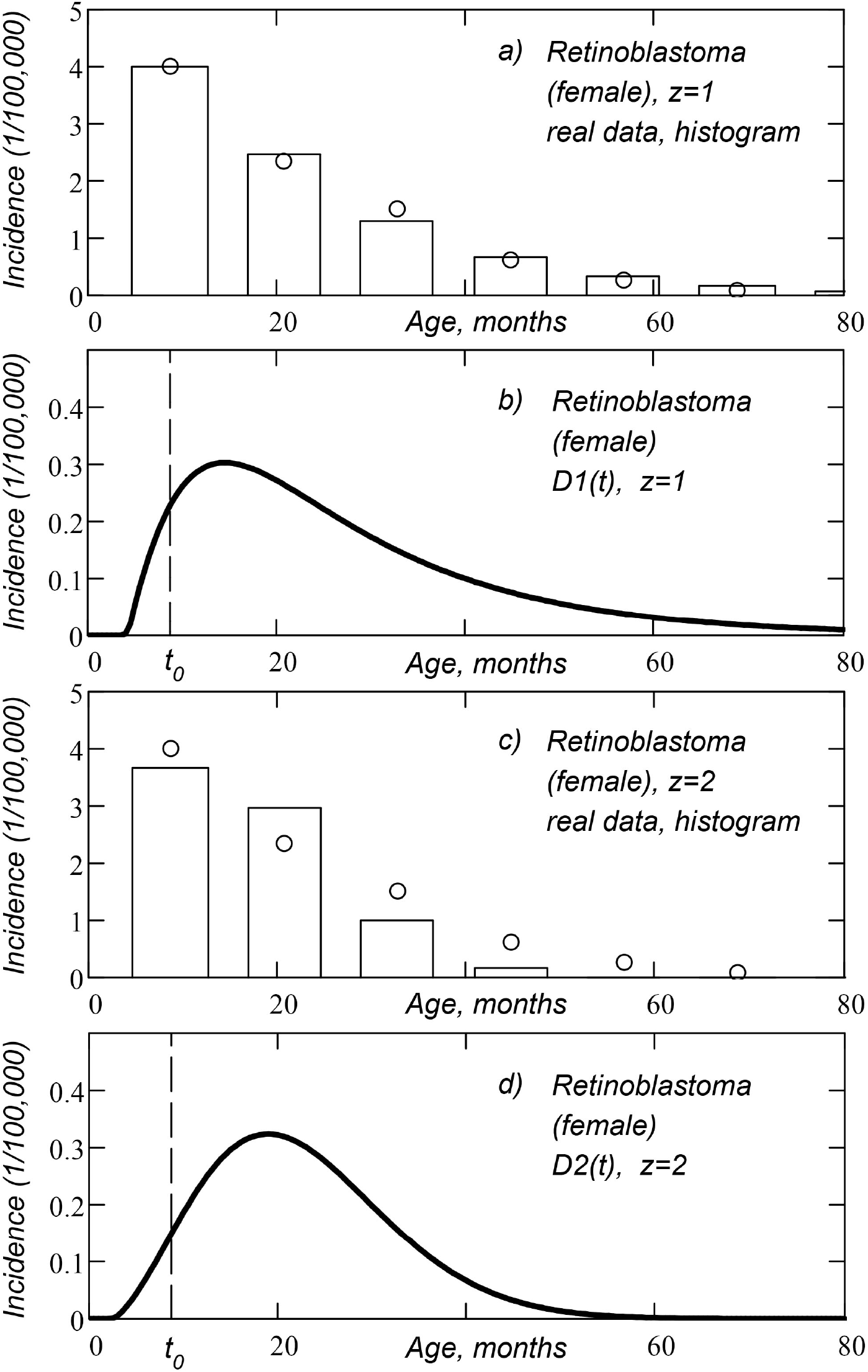
Age distribution of retinoblastomas, obtained for a complex mutational model (women): a) histogram of *D*1(*t*) function, real data set (open circles); b) *D*1(*t*) function of age distribution; c) histogram of *D*2(*t*) function, real data set; d) *D*2(*t*) function of age distribution. Here *z* is the number of cellular mutations required for the disease to occur. The vertical dashed line marks the time of birth *t* = *t*_0_.

**Figure 4.3.**
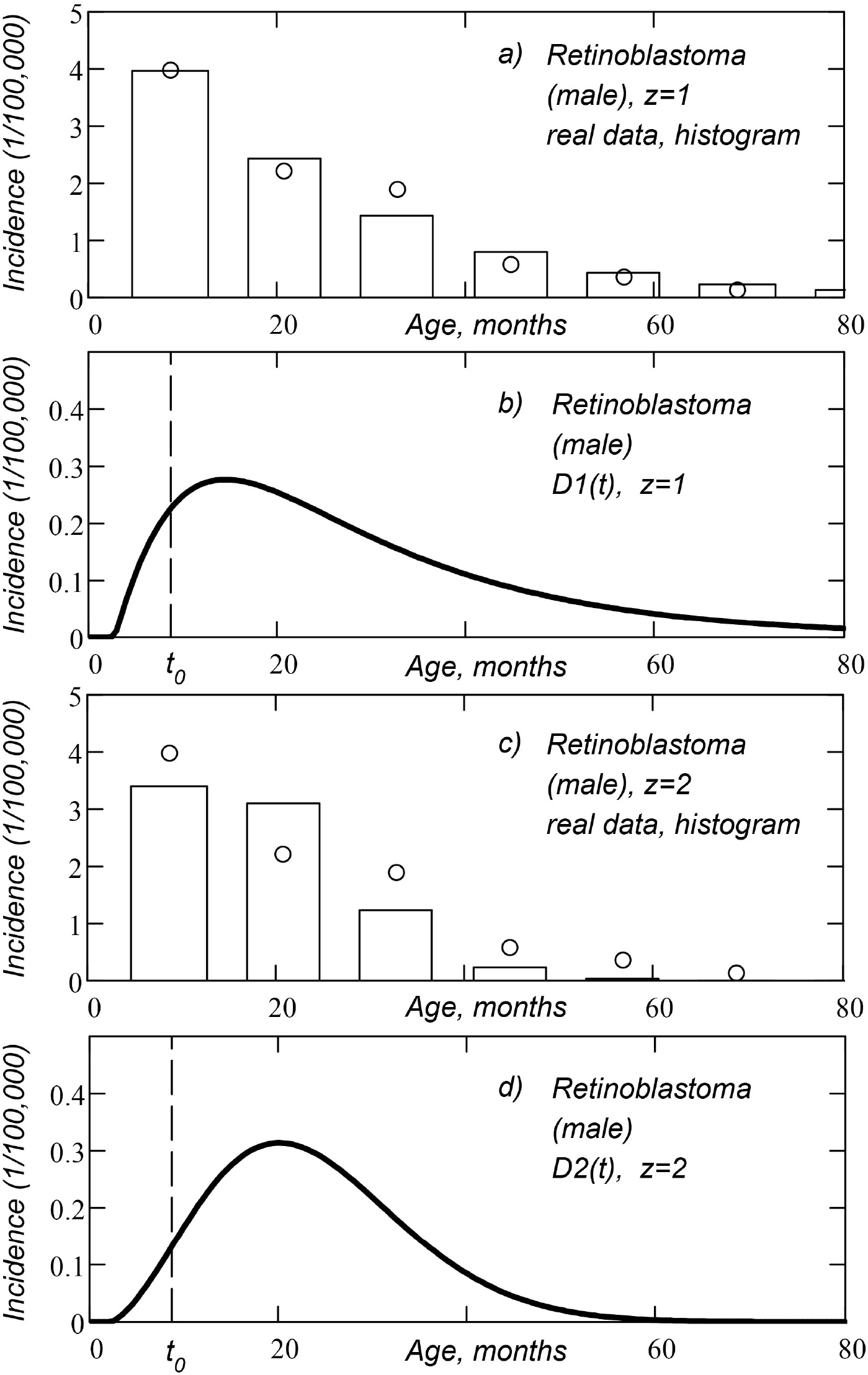
Age distribution of retinoblastomas, obtained for a complex mutational model (men): a) histogram of *D*1(*t*) function, real data set (open circles); b) *D*1(*t*) function of age distribution; c) histogram of *D*2(*t*) function, real data set; d) *D*2(*t*) function of age distribution. Here *z* is the number of cellular mutations required for the disease to occur. The vertical dashed line marks the time of birth *t* = *t*_0_.

**Figure 4.4.**
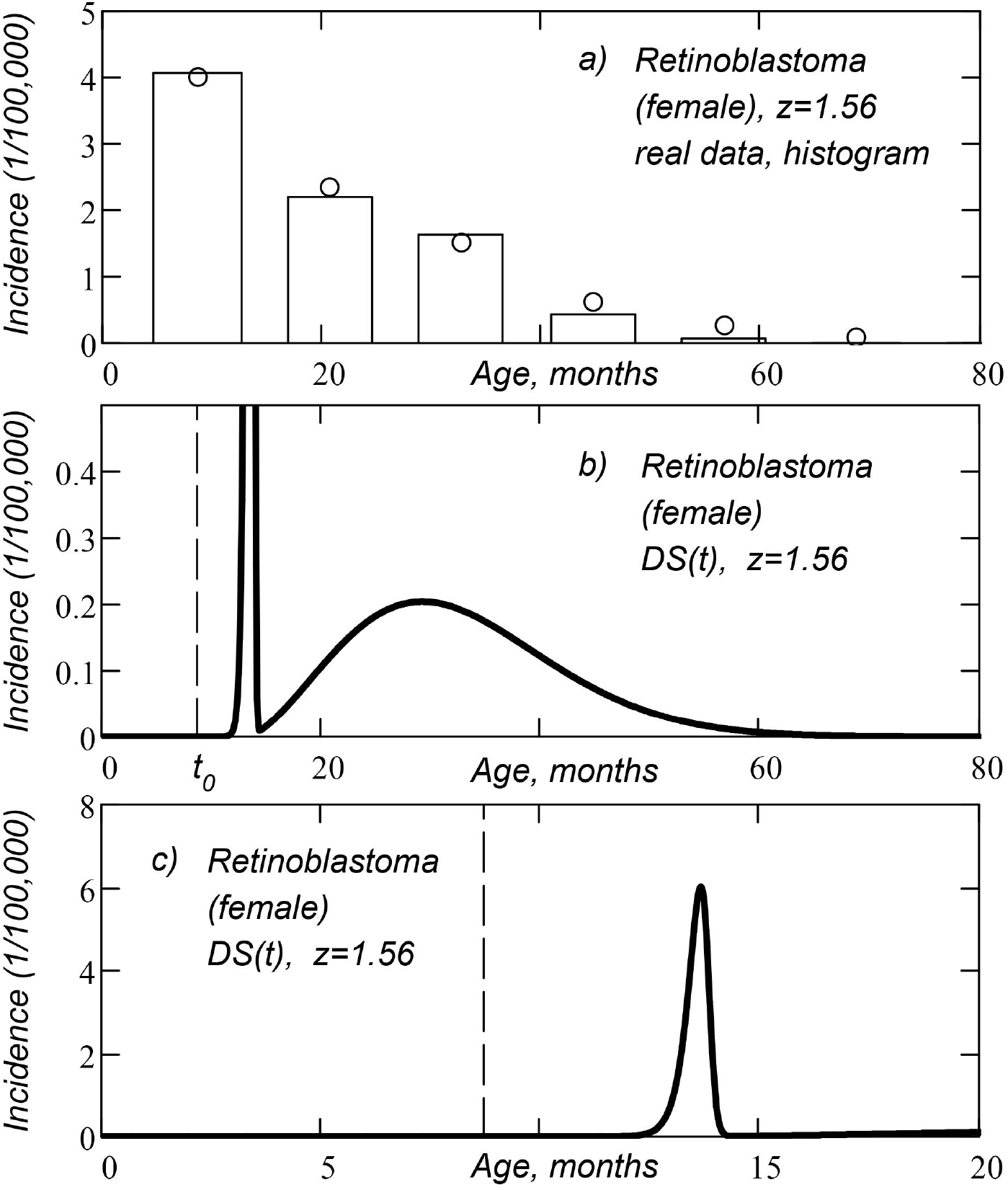
Composite age distribution of retinoblastomas, obtained for a complex mutational model (women): a) histogram of *DS*(*t*) function, real data set (open circles); b) *DS*(*t*) function of composite age distribution; c) *DS*(*t*) function of composite age distribution is presented in a different scale of the coordinate axes. Here *z* is the number of cellular mutations required for the disease to occur. The vertical dashed line marks the time of birth *t* = *t*_0_.

**Figure 4.5.**
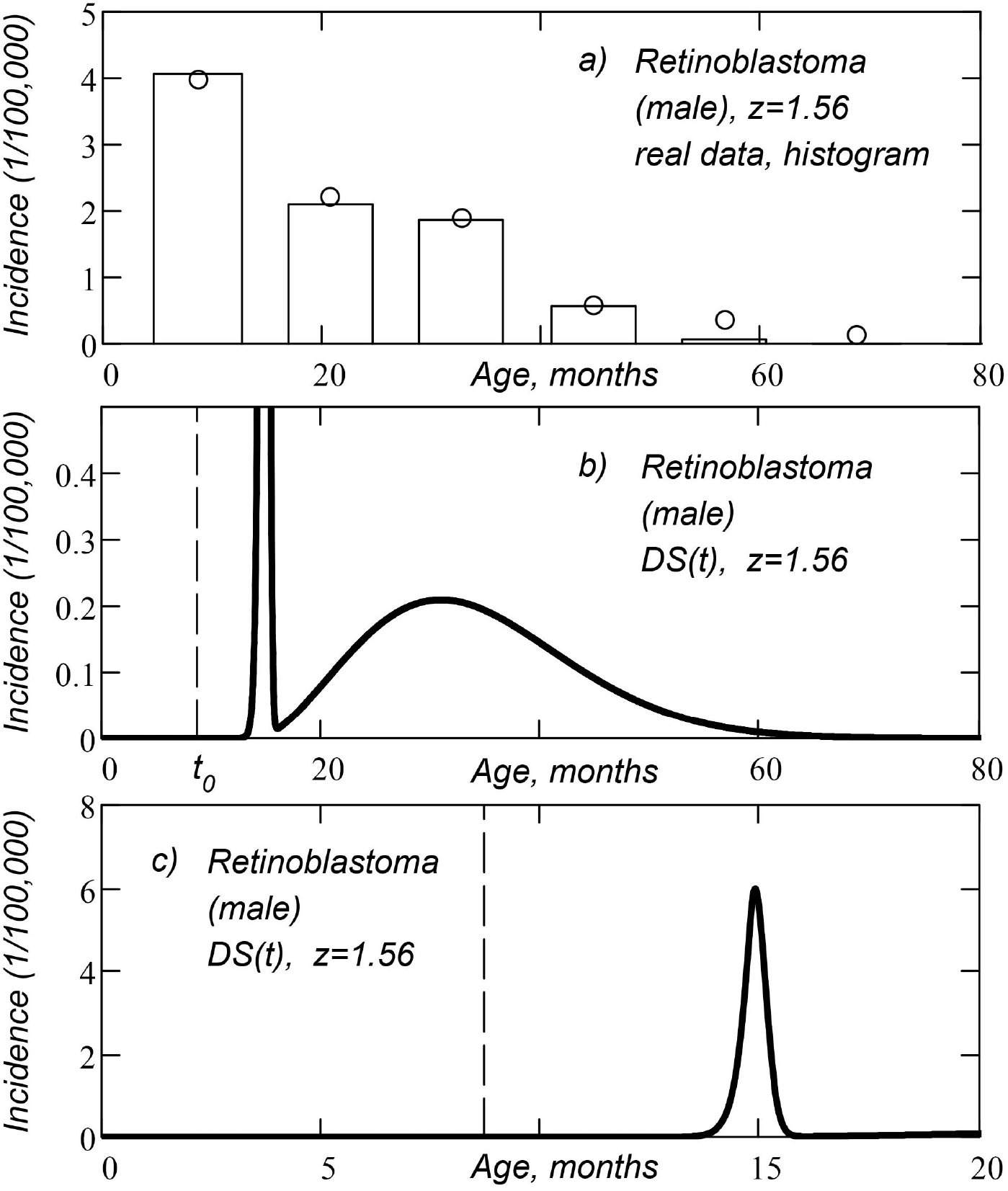
Composite age distribution of retinoblastomas, obtained for a complex mutational model (men): a) histogram of *DS*(*t*) function, real data set (open circles); b) *DS*(*t*) function of composite age distribution; c) *DS*(*t*) function of composite age distribution is presented in a different scale of the coordinate axes. Here *z* is the number of cellular mutations required for the disease to occur. The vertical dashed line marks the time of birth *t* = *t*_0_.

Real data in the figures are shown with open circles. Real data points are located along the time axis with one-year interval. The first point on the left in the figure corresponds to the time of birth (age is 0 years), the second point corresponds to the age of 1 year, and so on. The theoretical age distribution functions are shown with a bold curve line. The histograms plotted for the theoretical age distribution function are shown with rectangular bars. The vertical dashed line in the figures indicates the time of birth of the child *t* = *t*_0_.

Figures 4.4 and 4.5 show bimodal age distributions for which the incidence distribution function has two local maximums. The first peak of incidence (left maximum) corresponds to the group of patients with hereditary form of retinoblastoma, the second peak of incidence (right local maximum) corresponds to the group of patients with non-hereditary form of retinoblastoma.

An approximation for the case of two mutations (*z* = 2 in women and in men) gives a negative latency period (diagnosis lag time, parameter *T_S_* in Table 4.1). That is, if *T_S_* is negative, the value of the error function decreases. A negative TS parameter means that we can diagnose the disease before the key event that triggers the disease occurs. In practice, of course, this does not happen, so we write the value *T_S_* = 0 into the table.

For the compound age distribution (derived as the sum of hereditary and non-hereditary cases of retinoblastoma), we calculated the theoretical mean age at diagnosis (see Equations 4.12 and 4.13).

For hereditary female retinoblastoma, the mean age at diagnosis is: *t*_*d*1_ = 13.5 months; for non-hereditary female retinoblastoma, the mean age at diagnosis is: *t*_*d*2_ = 32.6 months;

For hereditary male retinoblastoma, the mean age at diagnosis is: *t*_*d*1_ = 14.9 months; for non-hereditary male retinoblastoma, the mean age at diagnosis is: *t*_*d*2_ = 34.2 months.

## 5. Third mutational model

This is the third mutational model in our works. The first two mutational models presented in [9] are called ‘‘simple mutational model” and ‘‘complex mutational model”. The simple model does not take into account the growth function of cell tissue and, for this reason, does not correspond to reality. The complex mutational model takes into account the increase in the number of cells in the cell tissue, therefore, it more accurately describes the process of obtaining of mutations. All mutational models assume that information about the obtaining mutations is transmitted from the parent cell to the daughter cells.

We have developed the third mutational model specifically for retinoblastoma. To avoid confusion in the names, we have named this model the “third mutational model”.

This model suggests that a key event in the cell damages only one allele of the Rb gene, and that cancer in the cell is activated only if two alleles of the Rb gene are damaged.

The third mutational model also suggests that the age distribution of retinoblastomas is formed by two different groups of patients. The first group has hereditary (inherited from one of the parents) Rb gene allele damage, and in order to initiate the disease, any individual retinal cell must receive only one mutation (allele damage). The second group of patients has two normal alleles in each cell, and in order to cause disease, the cell must receive two mutations (both alleles must be damaged).

As in the previous section, we consider the percentage of groups in the age distribution to be 44/56, based on the data presented in [16].

Thus, in contrast to the complex mutational model discussed above, in this model we believe that key events (mutations) occur not with the cell, but with one of the two alleles in the cell. That is, the a parameter in this model is the specific mutation frequency for one allele (the average number of key events that occur in one allele per unit time). Since in a group of healthy patients there are two normal alleles in the cell before the first mutation, the specific frequency of the first mutation should be twice as high. In the working equations, we assume that the frequency of the first mutation in these patients is 2a.

### 5.1. Age distribution function

In the third mutational model, the age distribution function *DS*(*t*) looks the same as the age distribution function in the complex mutational model:

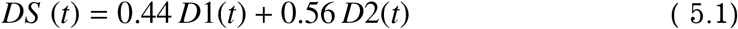

Here *D*1(*t*) and *D*2(*t*) functions are described by equations (4.1) and (4.2). The factors 0.44 and 0.56 correspond to the case when 44 percent of patients have hereditary retinoblastoma. If the reader has good statistics with a different percentage of these two patient groups, a different number (which corresponds to a different percentage) can be substituted into the equation.

The difference between the third mutational model and the complex mutational model is the equation for the function *N*1(*t*) (the number of cells with one or more mutations):

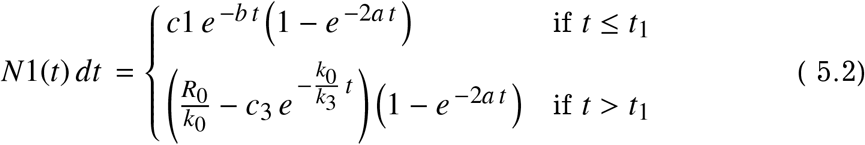

Note the double exponent power (before the last bracket in equations).

### 5.2. Approximation method

We approximate the real age distribution (Fig. 3.1) using the theoretical function (5.1). Function (5.1) corresponds to the case of a complex age distribution, when in 44% of patients (in the considered group of people) retinoblastoma occurs as a result of damage in one allele of Rb gene and in 56% of patients retinoblastoma occurs as a result of damage in both alleles.

Just as we did above, for the complex mutational model, we choose the optimal parameters *a, T_S_*, and *N_P_* for this function.

For each set of parameters (*a, T_S_*, and *N_P_*) in the age distribution function, we construct a histogram of the age distribution *DS*(*t*) (see Eq. 4.6-4.9) and calculate the error function (see Eq. 4.10). We believe that the optimal set of parameters gives the minimum value of the error function. The approximation is carried out programmatically, by the quasi-Newtonian method. For the obtained optimal data set, we also calculate the relative root-mean-square error (4.11).

### 5.3. Results

The approximation results are presented in Table 5.1 and Figures 5.1 and 5.2.

**Table 5.1.** Optimal parameters of approximation for the third mutational model.

| **Retinoblastoma** | $T_S$ | $a$ | $N_P$ | $Er$ | $Er\%$ |
| --- | --- | --- | --- | --- | --- |
| Female | 10.4 | $1.5 \cdot 10^{-5}$ | 8.4 | $4.79 \cdot 10^{-3}$ | 0.12 |
| Male | 11.8 | $1.5 \cdot 10^{-5}$ | 8.7 | $4.81 \cdot 10^{-3}$ | 0.12 |

**Figure 5.1.**
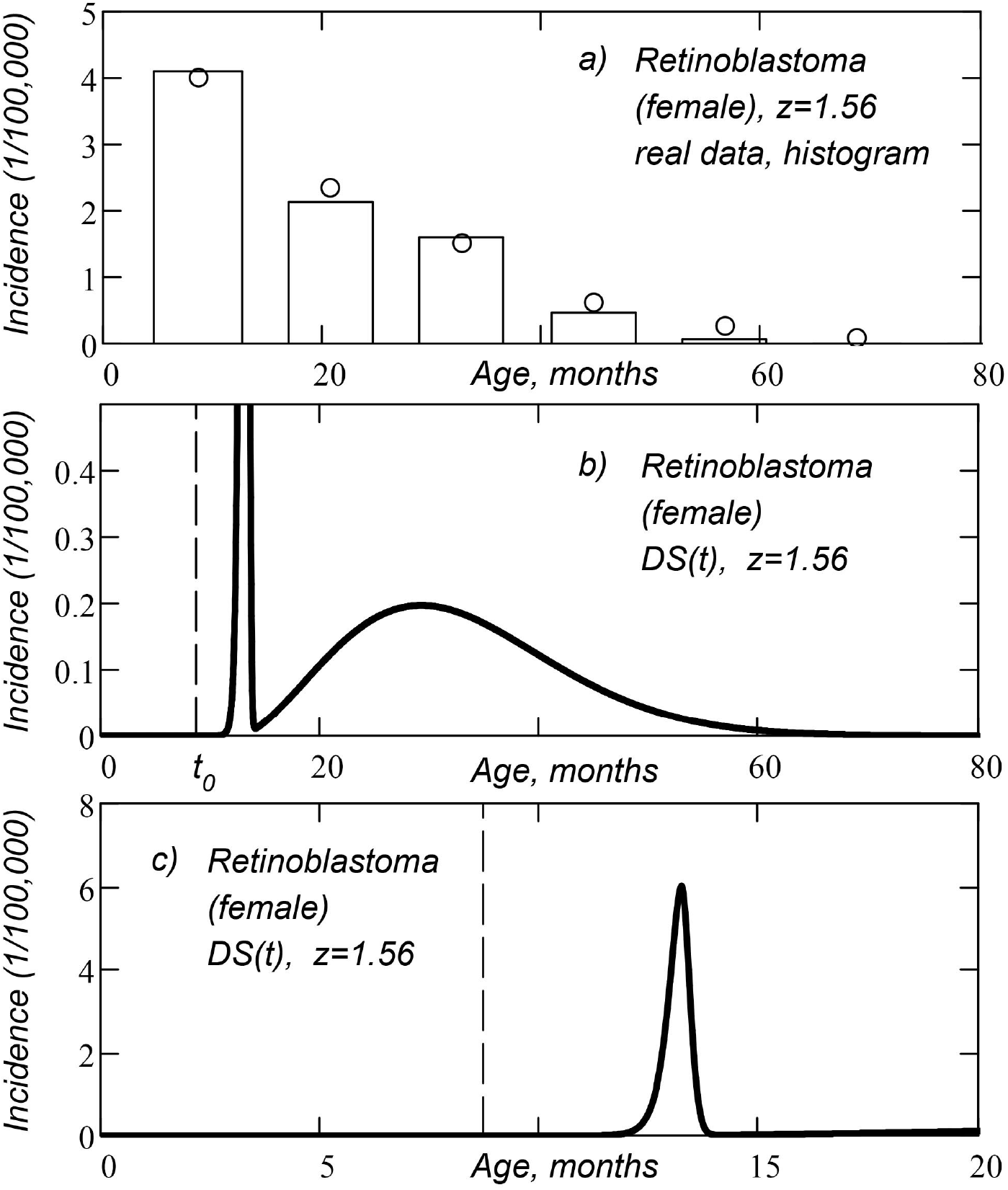
Composite age distribution of retinoblastomas, obtained for third mutational model (women): a) histogram of *DS*(*t*) function, real data set (open circles); b) *DS*(*t*) function of composite age distribution; c) *DS*(*t*) function of composite age distribution is presented in a different scale of the coordinate axes. Here *z* is the number of cellular mutations required for the disease to occur. The vertical dashed line marks the time of birth *t* = *t*_0_.

**Figure 5.2.**
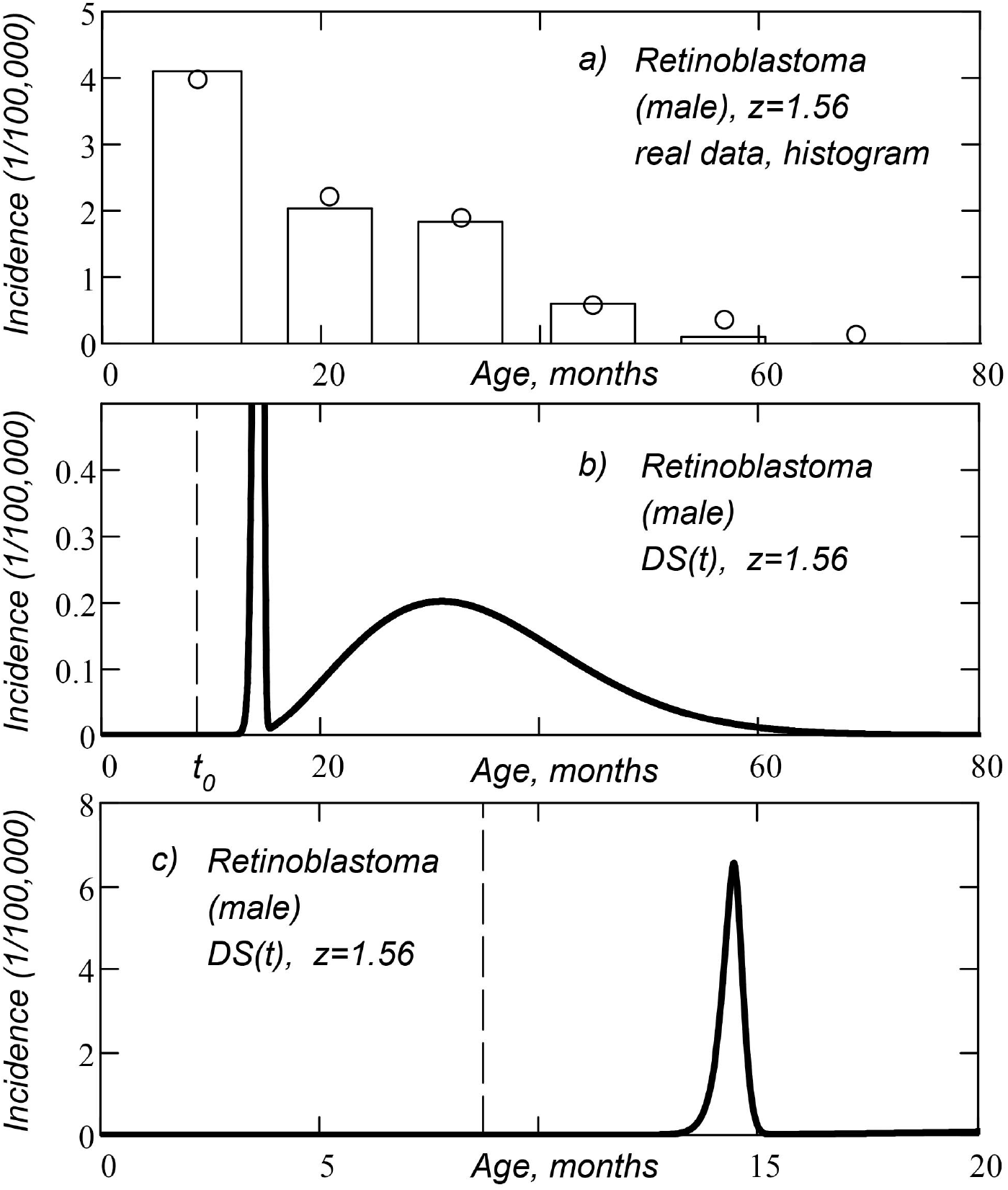
Composite age distribution of retinoblastomas, obtained for third mutational model (men): a) histogram of *DS*(*t*) function, real data set (open circles); b) *DS*(*t*) function of composite age distribution; c) *DS*(*t*) function of composite age distribution is presented in a different scale of the coordinate axes. Here *z* is the number of cellular mutations required for the disease to occur. The vertical dashed line marks the time of birth *t* = *t*_0_.

The *T_S_* parameter is the latent period of the disease (the time from the key event to the diagnosis).

The *a* parameter is the specific frequency of key events – the average number of key events that occur with one allele of Rb gene per unit of time (month).

The parameters *Er* and *Er_%_* are absolute and relative mean square error. The relative error is calculated as a percentage of the maximum value from the set of real data on the age distribution of retinoblastomas.

The obtained age distribution functions *DS*(*t*) with optimal approximation parameters, the histograms plotted for these functions, and real data sets are shown in Figures 5.1 (women) and 5.2 (men).

Real data in the figure are shown with open circles. Real data points are located along the time axis with an interval of 1 year.The first point on the left in the figure corresponds to the time of birth(age 0 years), the second point corresponds to the age of 1 year, and so on. The theoretical age distribution functions are shown by the bold curve. The histograms plotted for the theoretical age distribution function *DS*(*t*) are shown with rectangular bars. The vertical dashed line in the figures marks the time of birth *t* = *t*_0_.

For the resulting composite age distribution (the sum of hereditary and non-hereditary cases of retinoblastoma), we calculated theoretical mean age at diagnosis (see Eq. 4.12 and 4.13).

For hereditary female retinoblastoma, the mean age at diagnosis is: *t*_*d*1_ = 13.1 months; for non-hereditary female retinoblastoma, the mean age at diagnosis is:: *t*_*d*2_ = 32.8 months;

For hereditary male retinoblastoma, the mean age at diagnosis is: *t*_*d*1_ = 14.4 months; for non-hereditary male retinoblastoma, the mean age at diagnosis is:: *t*_*d*2_ = 34.4 months.

## 6. Retinoblastoma model with a sequence of key events

### 6.1. Mathematical model

In this model, we assume that a sequence of some (identical) key events that act on a group of cells leads to oncological disease. The number of cells in a group is *n_g_*.

In this model, instead of the cell tissue growth function *N*(*t*), we use the growth function of the number of cell groups *N_g_*(*t*). The *N_g_*(*t*) function is obtained by dividing the original cell tissue growth function by the number of cells in the group:

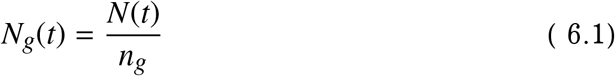

In this model, in contrast to the mutational model, information about key events (events that happened to the cell) is not transmitted from the parent cell to the daughter cells. Consequently, the equations describing the number of cells that have undergone a certain number of events (two, three, or more) are different from the equations for the mutation model. For the case of one key event, the equations are the same as for the mutation model with one mutation.

In this model, if a cell that has received a certain number of key events dies of old age, information about key events in the cell is lost, the number of cells with key events decreases, and these losses must be taken into account in the model equations.

In the case of retinoblastoma, we can ignore cell death from old age, and our equations will be relatively simple. It is assumed [5] that tumour cells of retinoblastoma originate from progenitor cells of the cones (light-sensitive cells of the retina of eye), and the cones live long enough. The lifespan of the cones is comparable to the lifespan of the human body.

First, we find the number of *N*0_*g*_ cell groups that have not received a single key event:

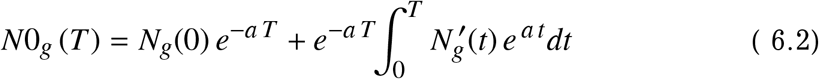

We obtained a similar formula in [9], when constructing a simple mutational model. In this case, a new term *N_g_*(0)*e*^-*aT*^ is added to the formula, since at the point *t* = 0 the function *N_g_*(*t*) is not equal to zero. Very often we can neglect this term, since at the initial moment of time the cell population consists of one cell, but in this case we include this term in the equation.

Here *a* is the specific frequency of key events (the average number of key events occurring with one cell group per unit time); *t* and *T* are the same time variable (different notation is used to avoid confusion with the limits of integration); 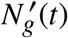 is the time derivative of the function *N_g_*(*t*).

The number of cell groups that received one key event (or more) is:

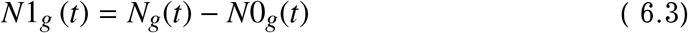

The number of cell groups that received exactly one key event (no more) is:

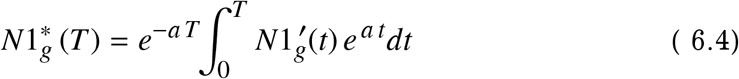

The number of cell groups that received two key events (or more) is:

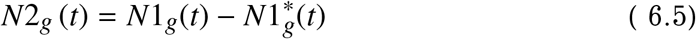

The number of cell groups that received exactly two key events (no more) is:

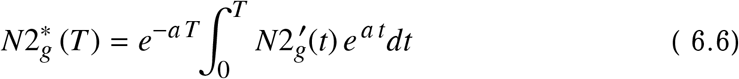

The number of cell groups that received three key events (or more) is:

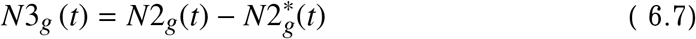

etc.

Acting in a similar way, we can obtain equations for functions describing the number of cell groups (in a given anatomical tissue) with any number of key events we need. We will need these formulas to construct the age distribution of oncological diseases.

For example, if we use equations (2.2) for the growth function of cell tissue, taking into account (6.2) we can write:

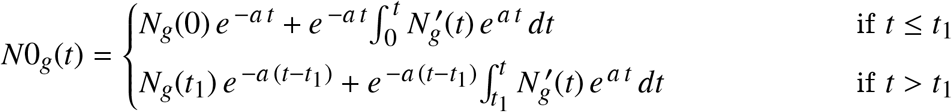

And *N*1_*g*_(*t*) function looks like this:

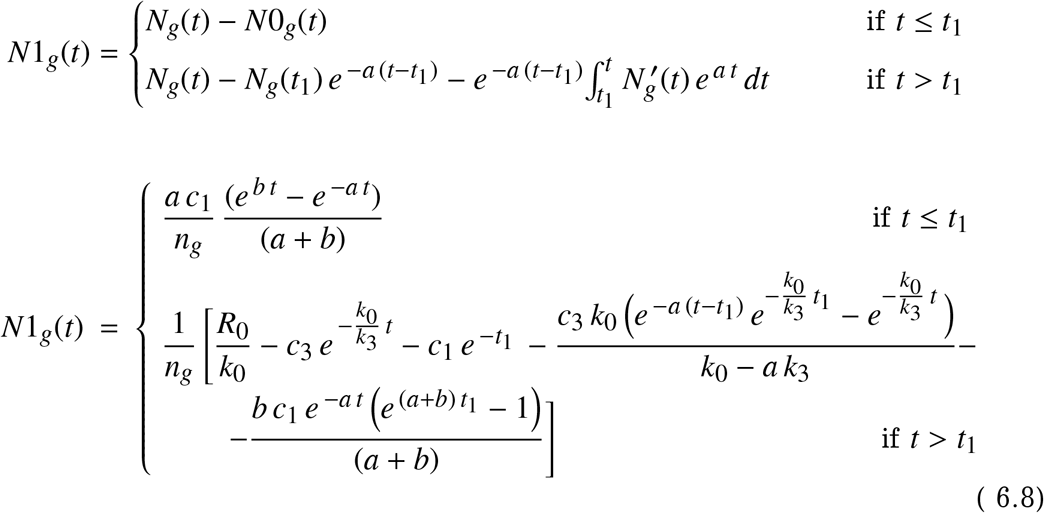

In the case of retinoblastoma, we will consider a sequence of no more than two key events, so we do not need *N*2_*g*_(*t*) and *N*3_*g*_(*t*) functions.

The functions *N_g_*(*t*), *N*0_*g*_(*t*), and *N*1_*g*_(*t*) are shown in Fig. 5.1 (in two scales of the coordinate axes). The time *t*_1_ is indicated in the figures with a vertical dashed line. When plotting the graphs, we used the following parameters: *c*_1_ = 1630.92; *R*_0_ = 5.08 · 10^5^; *k*_1_ = 1; = 0.058; *k*_3_ = 0.472; *n_g_* = 1; *t*_1_ = 2.87; *a* = 0.5.

As well as for the mutational model of retinoblastoma, we will consider the case when the age distribution of the incidence is formed by two groups of people: in the first group of patients for the onset of the disease, it is required that one key event occurs in the group of cells of anatomical tissue, in the second group of patients for the onset of the disease, two key events are required per group of cells.

Let the proportion of patients with one key event be 44 percent of the size of considered group (for which the age distribution is constructed); the proportion of patients with two key events is 56 percent. That is, the percentage of groups is the same as we used earlier for the mutational model.

This ratio defines the formula for the compound age distribution (see below).

**Figure 6.1.**
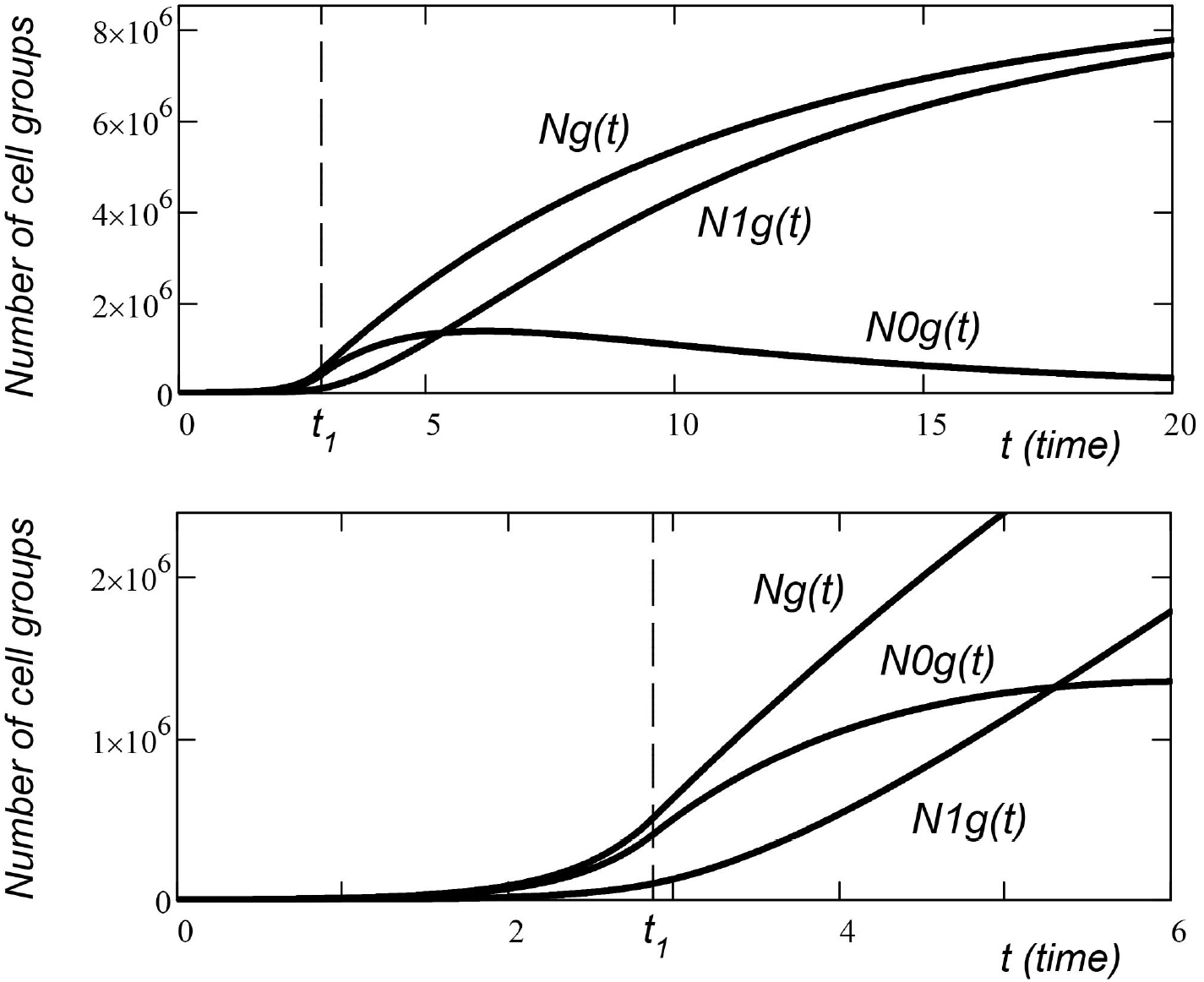
Functions: *N_g_*(*t*); *N*0_*g*_(*t*); *N*1_*g*_(*t*) (the number of cell groups in the cell tissue; the number of cell groups that did not receive key events; the number of cell groups that have one or more key events). Identical functions are shown in the upper and bottom figures at different scales of the coordinate axes.

### 6.2. Age distribution functions

Using the equations obtained in [9], we can write down the functions *D*1(*t*) and *D*2(*t*), which describe the age distributions of retinoblastomas. The *D*1(*t*) function corresponds to the case when a disease in a cell group occurs after the cell group receives one key event. The *D*2(*t*) function corresponds to the case when a disease in a group of cells occurs after two key events.

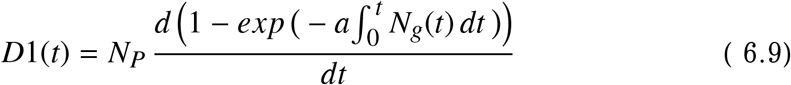

Here the constant *N_P_* is the number of people in the group for which the age distribution is constructed; *N_g_*(*t*) is a function that describes the growth of the number of cell groups in the anatomical tissue.

It should be remembered that the growth function (in our description) is the piecewise function *N_g_*(*t*), which is given by different equations at two different time intervals (before *t*_1_ and after *t*_1_). Therefore, after the time *t*_1_, the integration should be carried out separately for each time interval:

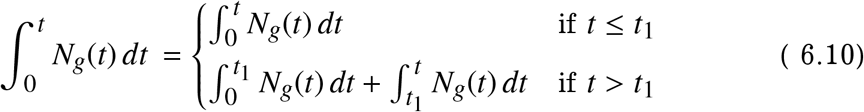

The function *D*2(*t*) (age distribution for the case when the disease occurs after two key events in a group of cells) is as follows:

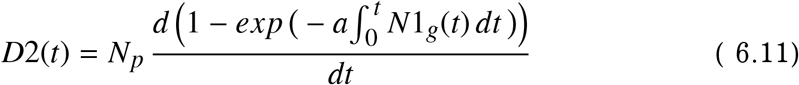

Also, do not forget about the integration of the piecewise function:

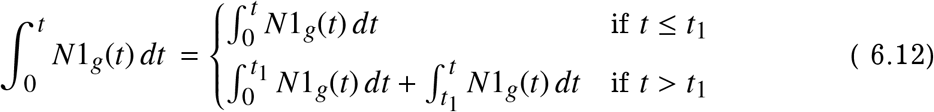

The composite age distribution *DS*(*t*) (constructed for two functions *D*1(*t*) and *D*2(*t*), taken in the ratio 44/56) is:

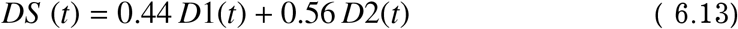

Examples of the function of the composite age distribution *DS*(*t*) are shown in Fig. 6.2. The figure shows the functions for a different number of cells in a cell group (a group of cells that is affected by a common oncogenic event). When plotting the graphs, we used the following parameters: *R*_0_ = 2.78 · 10^5^, *k*_0_ = 10.30, *k*_1_ = 1, *k*_3_ = 10.22, *k* = 5.87 · 10^5^, *t*_1_ = 2.8, *a* = 10.003, *N_p_* = 8.3.

**Figure 6.2.**
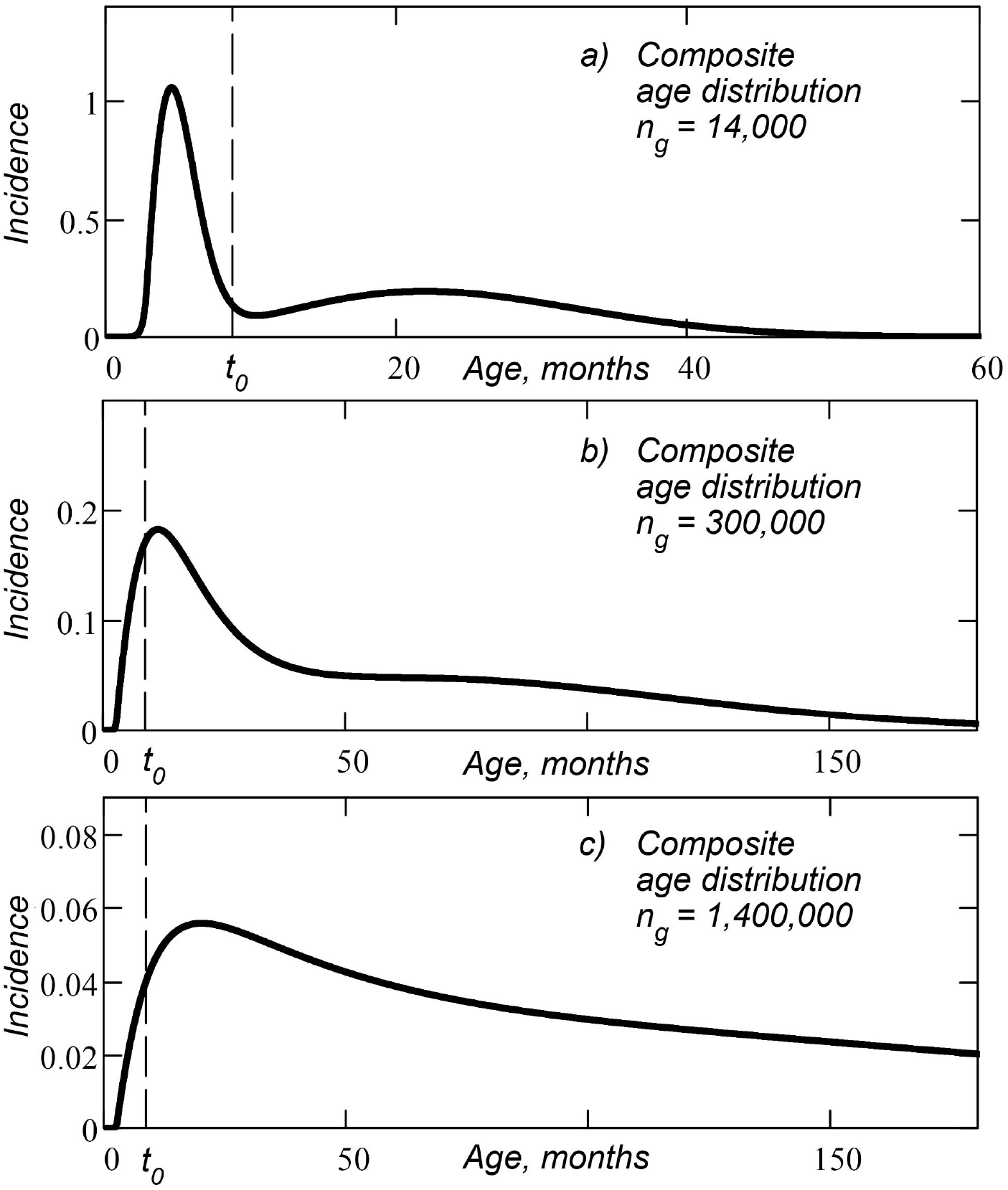
Composite age distribution of retinoblastomas, obtained for the model with a sequence of key events for different *n_g_* parameter (the number of cells in a cell group affected by a common oncogenic event). The vertical dashed line marks the time of birth *t* = *t*_0_.

### 6.3. Approximation method

The approximation of the real age distribution (Figure 3.1) is carried out using the theoretical functions (6.10), (6.12), (6.14). Function (6.10) corresponds to the case when retinoblastoma occurs after one carcinogenic event (which happened to a group of cells). Function (6.12) corresponds to the case when retinoblastoma occurs after two carcinogenic events (which happened to one group of cells). Function (6.12) corresponds to the composite case, when in 44 percent of patients (from the considered group of people) retinoblastoma occurs after one carcinogenic event (that happened to a group of cells), in 56 percent of patients retinoblastoma occurs after two carcinogenic events (that happened to one group of cells).

Just as we did it for the mutational model, we choose the optimal parameters *a, T_S_*, and *N_P_* for these functions.

Also in this model we optimize the additional parameter *n_g_* (number of cells in the group where the carcinogenic event occurs).

For the case of one key event (*z* = 1), the parameters *a* and *n_g_* enter the equations as the ratio *a*/*n_g_*. When approximating, the change in the *a* parameter can be compensated by the change in the *n* parameter, and we cannot separate these parameters (we cannot calculate these parameters separately, but we can only calculate the ratio *a*/*n_g_*). Therefore, only for this case, we consider that the parameter *n_g_* is equal to one.

For *n_g_* = 1, the equations for *D*1(*t*) function are identical to the equations for the case of one key event in the complex mutational model (for which we optimized the parameters in the previous section). Therefore, the data in the results table for the case *z* = 1 (one key event) coincides with the data in table 4.1 (first line).

As well as for the mutational model, for each set of parameters *n_g_, a, T_S_* and *N_P_* in the age distribution function, we construct a histogram of the age distribution (see formulas 4.6-4.9) and calculate the error function for it:

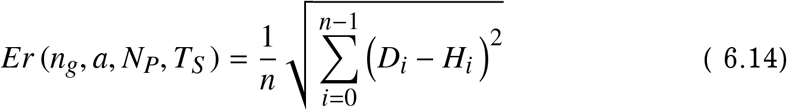

We believe that the optimal set of parameters gives the minimum value of the error function. The approximation is carried out programmatically, by the quasi-Newtonian method.

For the obtained optimal data set, we also calculate the relative root-meansquare error:

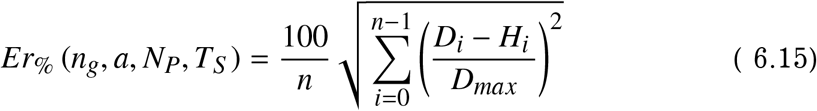

The function shows the relative root mean square error of approximation as a percentage of the maximum value of *D_max_* from a set of real data.

### 6.4. Results

The approximation results are presented in Table 6.1.

**Table 6.1.** Optimal approximation parameters for the model with sequence of key events.

| **Retinoblastoma** | $n_g$ | $T_S$ | $a$ | $N_P$ | $Er$ | $Er\%$ |
| --- | --- | --- | --- | --- | --- | --- |
| Female; z=1 | 1 | 1.6 | $6.3 \cdot 10^{-9}$ | 9.1 | $3.41 \cdot 10^{-3}$ | 0.09 |
| Female; z=1.56 | $7.5 \cdot 10^5$ | 3.6 | $1.8 \cdot 10^{-2}$ | 8.8 | $1.83 \cdot 10^{-3}$ | 0.05 |
| Female; z=2 | $9.2 \cdot 10^6$ | 0 | $1.1 \cdot 10^{-1}$ | 8.2 | $9.92 \cdot 10^{-3}$ | 0.25 |
| Male; z=1 | 1 | 0.4 | $5.4 \cdot 10^{-9}$ | 9.6 | $7.18 \cdot 10^{-3}$ | 0.18 |
| Male; z=1.56 | $1.3 \cdot 10^5$ | 8.5 | $8.5 \cdot 10^{-3}$ | 8.9 | $3.98 \cdot 10^{-3}$ | 0.12 |
| Male; z=2 | $9.2 \cdot 10^6$ | 0 | $1.1 \cdot 10^{-1}$ | 8.2 | $13.94 \cdot 10^{-3}$ | 0.35 |

The *z* parameter is the number of key events (in a group of cells) that initiate the disease. The parameter *z* = 1.56 refers to a composite case (44 percent of the group are patients with one mutation, 56 percent are patients with two mutations).

The *n_g_* parameter is the number of cells in one cell group (the group affected by the common carcinogenic event).

The *T_S_* parameter is the latency period of the disease (time from the key event that initiates the cancer to the diagnosis).

The *a* parameter is the specific frequency of key events – the average number of key events that happen to a group of cells per unit of time (month).

The parameters *Er* and *Er_%_* are absolute and relative mean square error. the maximum value from the set of real data on the age distribution of retinoblastomas.

The first lines in Tables 6.1 (for women and men) coincide with the first lines in Tables 4.1, because for one key event (*z* = 1) with *n_g_* = 1, the theoretical equations for this model coincide with the equations for the mutational model with one mutation.

The obtained age distribution functions with optimal approximation parameters, histograms constructed for these functions, as well as real data sets are shown in Figures 6.2, 6.3, 6.4, and 6.5.

**Figure 6.3.**
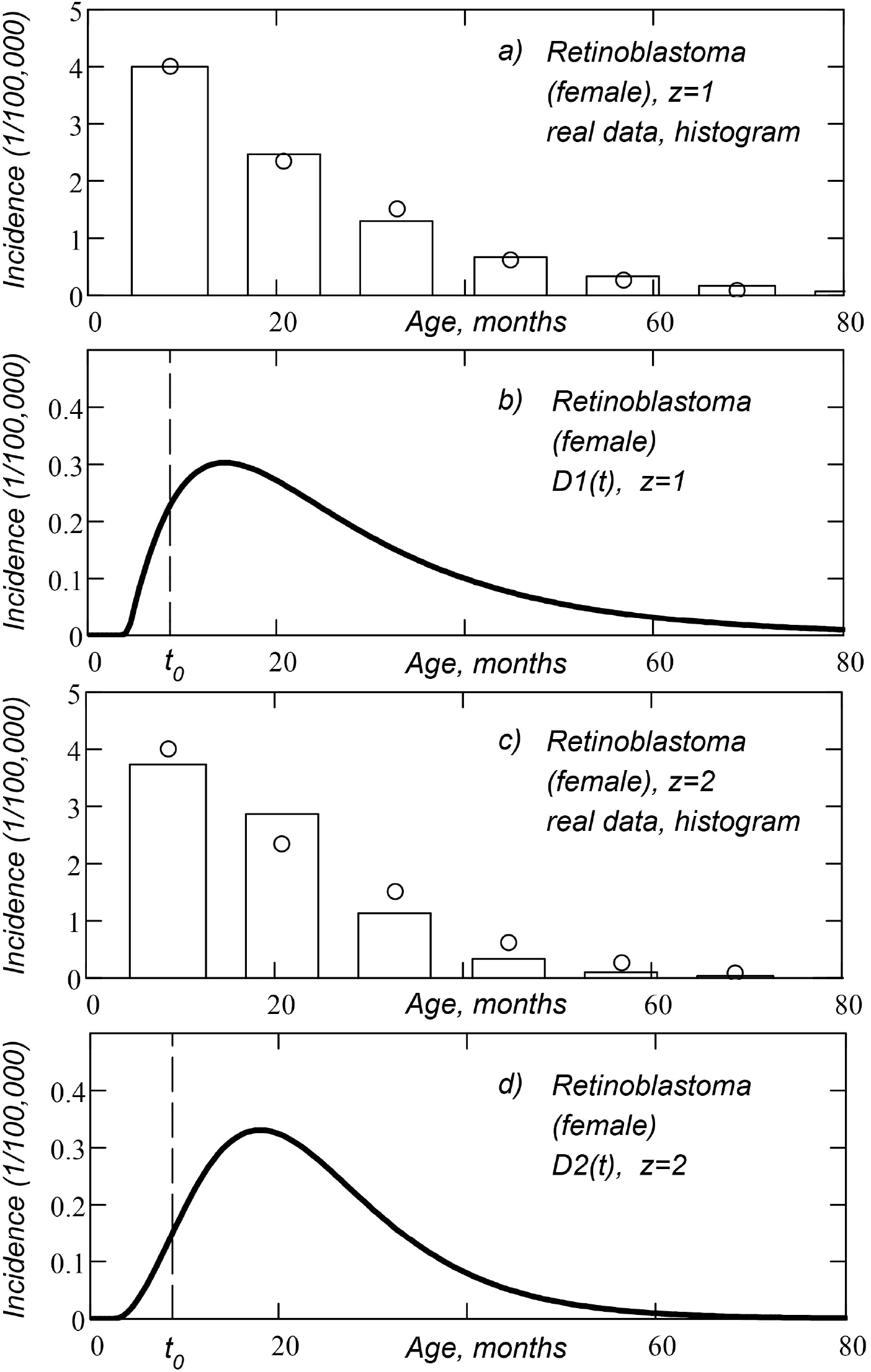
Age distribution of retinoblastomas, obtained for the model with a sequence of key events (women): a) histogram of D1(t) function, real data set (open circles); b) *D*1(*t*) function of age distribution; c) histogram of *D*2(*t*) function, real data set (open circles); d) *D*2(*t*) function of age distribution. Here *z* is the number of mutations in one cell group required for the cancer to occur. The vertical dashed line marks the time of birth *t* = *t*_0_.

**Figure 6.4.**
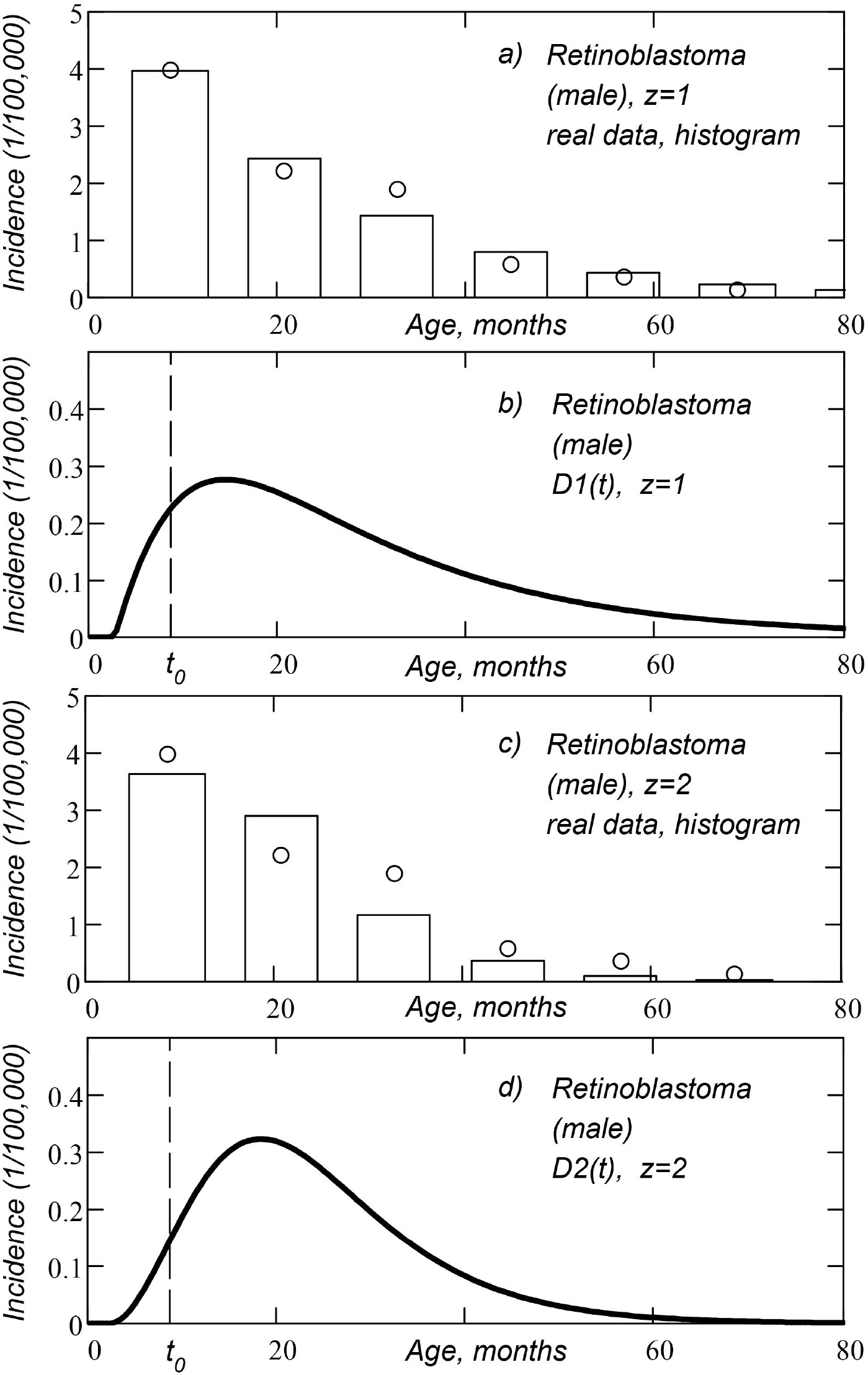
Age distribution of retinoblastomas, obtained for the model with a sequence of key events (men): a) histogram of *D*1(*t*) function, real data set (open circles); b) *D*1(*t*) function of age distribution; c) histogram of *D*2(*t*) function, real data set (open circles); d) *D*2(*t*) function of age distribution. Here *z* is the number of mutations in one cell group required for the cancer to occur. The vertical dashed line marks the time of birth *t* = *t*_0_.

**Figure 6.5.**
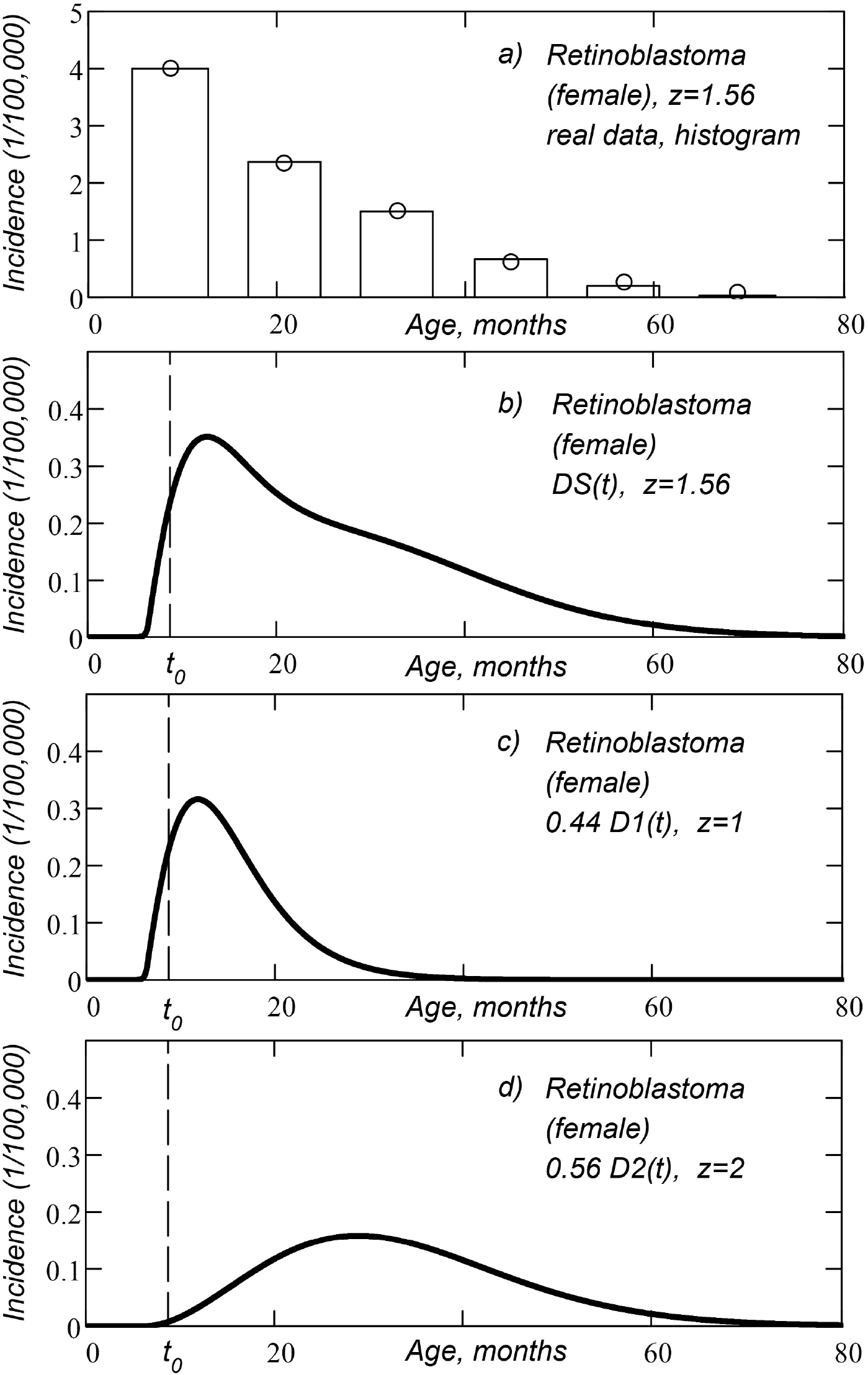
Composite age distribution of retinoblastomas, obtained for the model with a sequence of key events (women): a) histogram of *DS*(*t*) function, real data set (open circles); b) *DS*(*t*) function of composite age distribution; c) *D*1(*t*) function of age distribution (hereditary retinoblastoma). d) *D*2(*t*) function of age distribution (non-hereditary retinoblastoma). Here *z* is the number of key events in a group of cells required for cancer to occur. The vertical dashed line marks the time of birth *t* = *t*_0_.

**Figure 6.6.**
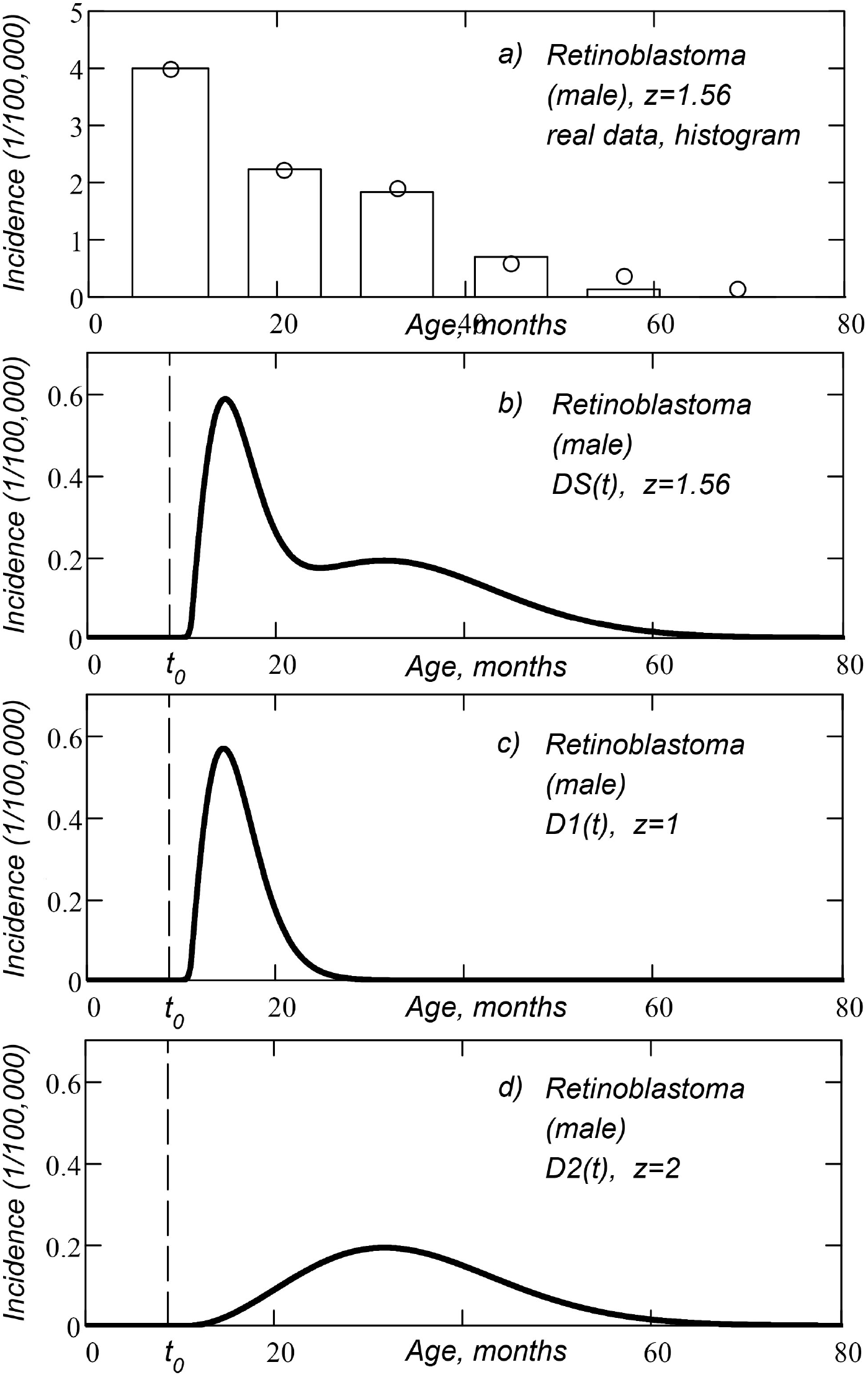
Composite age distribution of retinoblastomas, obtained for the model with a sequence of key events (men): a) histogram of *DS*(*t*) function, real data set (open circles); b) *DS*(*t*) function of composite age distribution; c) *D*1(*t*) function of age distribution (hereditary retinoblastoma). d) *D*2(*t*) function of age distribution (non-hereditary retinoblastoma). Here *z* is the number of key events in a group of cells required for cancer to occur. The vertical dashed line marks the time of birth *t* = *t*_0_.

Real data in the figures are shown by open circles. Real data points are located along the time axis with an interval of 1 year. The first point on the left in the figure corresponds to the time of birth (age 0 years), the second point corresponds to the age of 1 year, and so on. The theoretical age distribution functions are shown by the bold curve. The histograms plotted for the theoretical age distribution function are shown with rectangular bars. The vertical dashed line in the figures marks the time of birth *t* = *t*_0_.

For the resulting composite age distributions (the sum of hereditary and non-hereditary cases of retinoblastoma), we calculated theoretical mean age at diagnosis (see Eq. 4.12 and 4.13).

For hereditary female retinoblastoma, the mean age at diagnosis is: *t*_*d*1_ = 14.8 months; for non-hereditary female retinoblastoma, the mean age at diagnosis is: *t*_*d*2_ = 33.6 months.

For hereditary male retinoblastoma, the mean age at diagnosis is: *t*_*d*1_ = 16.ø months; for non-hereditary male retinoblastoma, the mean age at diagnosis is: *t*_*d*2_ = 34.7 months.

## 7. Model of single carcinogenic event with two different latencies

To have an objective understanding of the mechanisms of development of retinoblastoma, we must also consider a disease model for the case of one key event, but with two different groups of patients, each of which has a different latency period of the disease.

It can be assumed that in patients with bad heredity (genes obtained from a parent who had retinoblastoma) the disease develops faster (short latency period), and, on the contrary, in patients with good heredity the disease (which arose also accidentally, as in the first group) develops slower. (longer delay period).

In this section we consider such a model of retinoblastoma.

### 7.1. Age distribution function

In this oncological model, we assume that the oncological disease is initiated by one key event, but in two different groups of patients the disease has a different latency period (the time between the initiating key event and the diagnosis of the disease). One group (44 percent of the group of people for which the age distribution is constructed) has a latency period *T*_*S*1_, the other group (56 percent) has a latency period *T*_*S*2_. Here we assume that *T*_*S*1_ < *T*_*S*2_.

As mentioned earlier, we selected this percentage based on the data presented in [16] (918 hereditary and non-hereditary cases of retinoblastoma).

The function *DS*(*t*) of the composite age distribution for this model has the following form:

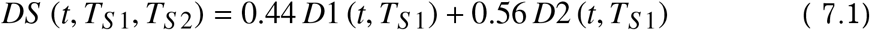

The compound age distribution function depends on the time t and on two parameters: *T*_*S*1_ and *T*_*S*2_. The function *D*1 (*t*, *T*_*S*1_) is the age distribution *D*1(*t*), shifted to the right along the horizontal time axis by *T*_*S*1_. The function *D*1(*t*) is given by equation (6.10), which we used earlier for the case when a cancer is initiated by one key event in a cell or group of cells.

Similarly, the function *D*1 (*t*, *T*_*S*2_) is the function *D*1(*t*), shifted to the right along the time axis by *T*_*S*2_.

### 7.2. Approximation method

The approximation of the real age distribution (Fig. 3.1) is carried out using the theoretical function (7.1).

We optimize the function parameters: *a*, *T*_*S*1_, *T*_*S*2_, and *N_P_*. Here *a* is the specific frequency of key events (the average number of key events that happen to a group of cells per unit of time); *T*_*S*1_ and *T*_*S*2_ are disease latencies for two groups of patients (hereditary and non-hereditary retinoblastoma); *N_P_* is the number of people in the group for which the age distribution of retinoblastomas is constructed.

When approximating the data, we assume that the parameter *n_g_* (the number of cells in the group with which a carcinogenic event occurs) is equal to one, because, for the case when cancer is initiated by one key event, the parameters a and *n_g_* always enter the equations in the form of the ratio *a*/*n_g_*, and an increase in one parameter can be exactly compensated by an increase in another parameter. Both parameters are currently unknown to scientists, since no practical measurements of these parameters have been carried out.

As before, for each set of parameters: a, *T*_*S*1_, *T*_*S*2_, and *N_P_* in the age distribution function, we construct a histogram of the age distribution *DS*(*t*) (see Eq. 4.6-4.9) and calculate the error function *Er* (least squares method):

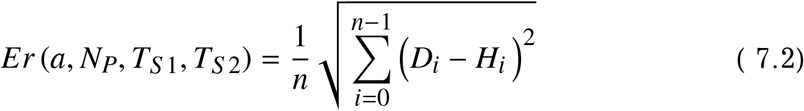

Here *D_i_* is the *i*-th value from the real age distribution; *H_i_* is the value of the *i*-th bar of the histogram built for the theoretical function *DS*(*t*) of the age distribution; n is the number of points in the real dataset.

We believe that the optimal set of parameters gives the minimum value of the error function. The approximation is carried out programmatically, by the quasi-Newtonian method. For the obtained optimal dataset, we also compute the relative root-mean-square error *Er*_%_:

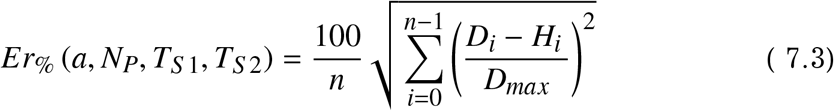

The function shows the relative root mean square error of approximation as a percentage of the maximum value of *D_max_* from a set of real age distribution data (the real age distribution is shown in Fig. 3.1).

### 7.3. Results

The approximation results are presented in table 7.1.

**Table 7.1.** Optimal approximation parameters for a model of a single carcinogenic event with two different latencies.

| **Retinoblastoma** | $T_{S1}$ | $T_{S2}$ | $a$ | $N_P$ | $Er$ | $Er\%$ |
| --- | --- | --- | --- | --- | --- | --- |
| Female; $z=1$ | 0.6 | 1.0 | $6.1 \cdot 10^{-9}$ | 9.1 | $3.24 \cdot 10^{-3}$ | 0.08 |
| Male; $z=1$ | 0.1 | 0.3 | $5.4 \cdot 10^{-9}$ | 9.5 | $7.17 \cdot 10^{-3}$ | 0.18 |

In this case, the number of key events (in a cell or group of cells) initiating a disease is equal to one (parameter *z* = 1).

The *T*_*S*1_ parameter is the latent period of the disease (the time from the key event initiating the disease to the diagnosis) in a group of patients with hereditary retinoblastoma. The *T*_*S*2_ parameter is the latent period of the disease in a group of patients with a non-hereditary form of retinoblastoma.

The *a* parameter is the specific frequency of key events – the average number of key events that happen to a cell or a group of cells per unit of time (month).

The parameters *Er* and *Er_%_* are the absolute and relative root mean square error. The relative error is calculated as a percentage of the maximum value from the set of real data on the age distribution of retinoblastomas.

The obtained age distribution functions with optimal approximation parameters, histogram plotted for these functions, as well as real data sets are shown in Figure 7.1.

**Figure 7.1.**
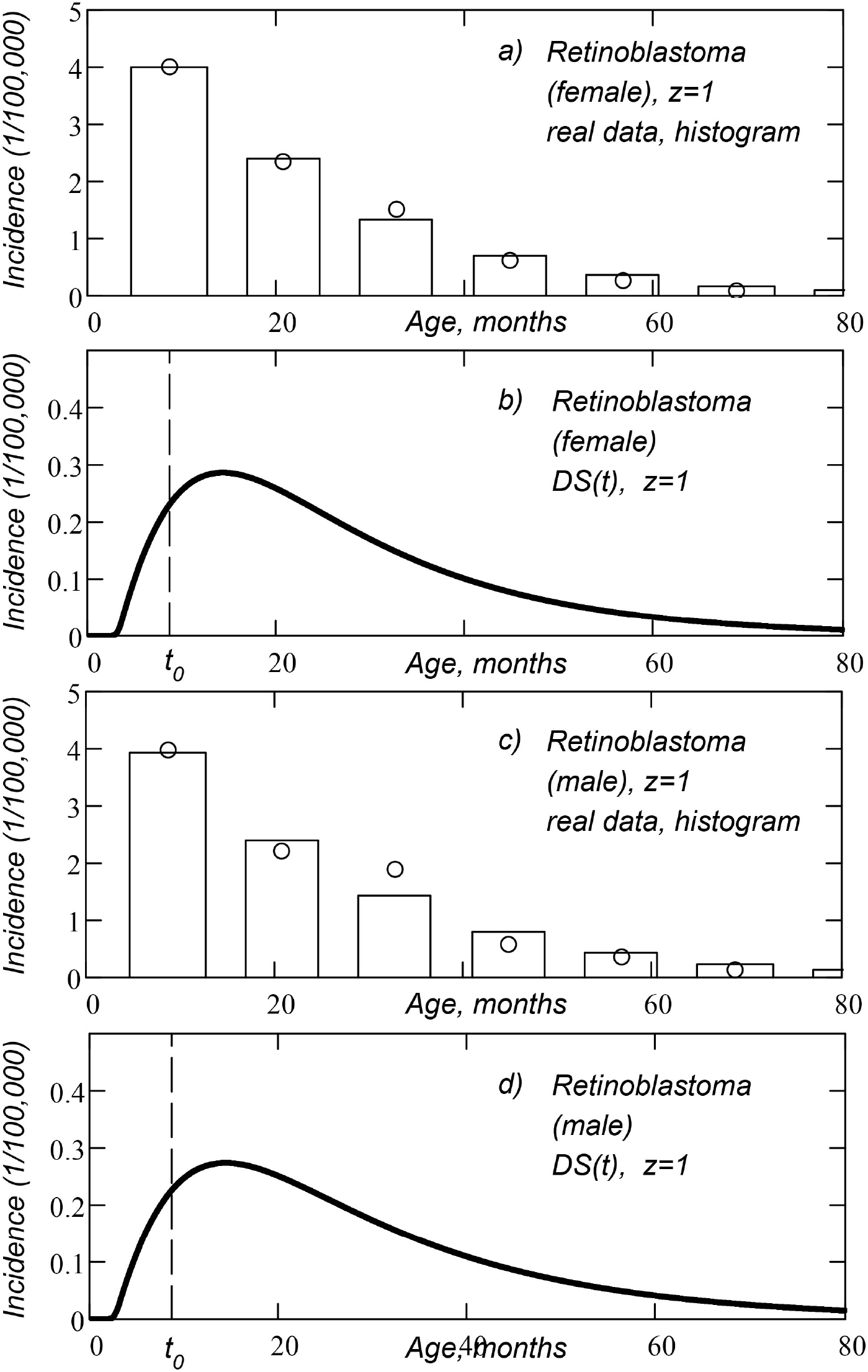
Composite age distribution of retinoblastomas, obtained for the model of single carcinogenic event with two different latencies: a) histogram of *DS*(*t*) function (women), real data set (open circles); b) *DS*(*t*) function of composite age distribution (women); c) histogram of *DS*(*t*) function (men), real data set (open circles); d) *DS*(*t*) function of composite age distribution (men). The vertical dashed line marks the time of birth *t* = *t*_0_.

Real data in the figures are shown with open circles. Real data points are located along the time axis with an interval of 1 year. The first point on the left in the figure corresponds to the time of birth (age 0 years), the second point corresponds to the age of 1 year, and so on. The theoretical age distribution functions are shown by the bold curve. The histograms plotted for the theoretical age distribution function are shown with rectangular bars. The vertical dashed line in the figures marks the time of birth *t* = *t*_0_.

The age distribution function in Fig. 7.1 has one maximum, although it is a composite distribution that is the sum of two distributions for two patient groups (hereditary and non-hereditary retinoblastoma). The highs (tops) of two different distributions are close to each other and merge into one common maximum.

For the resulting composite age distribution (sum of inherited and noninherited cases of retinoblastoma), we calculated the theoretical mean age at diagnosis (see Eq. 4.12 and 4.13).

For hereditary female retinoblastoma, the mean age at diagnosis is: *t*_*d*1_ = 25.9 months; for non-hereditary female retinoblastoma, the mean age at diagnosis is: *t*_*d*2_ = 26.5 months;

For hereditary male retinoblastoma, the mean age at diagnosis is: *t*_*d*1_ = 27.5 months; for non-hereditary male retinoblastoma, the mean age at diagnosis is: *t*_*d*2_ = 28.2 months.

## 8. Discussion

The approximation of real data, performed in this article, makes it possible to determine the following parameters of retinoblastoma:

– the number of key events (mutations) per cell (per one allele of Rb gene for the third mutational model) required for cancer to occur (*z* parameter);
– the average specific frequency of key events (mutations) per unit time (month) per cell (per one allele of Rb gene for the third mutational model) in the considered cell tissue (this is a parameter in the presented tabular results);
– the latent period of the disease (time interval between the appearance of the first cancer cell and detection of cancer) (*T_S_* parameter in the presented tabular results);
– theoretical mean age at diagnosis (for compound age distributions) for hereditary and non-hereditary forms of retinoblastoma (parameters *t*_*d*1_ and *t*_*d*2_);

The *N_P_* parameter in the table shows what proportion of people from the considered group develop cancer (during their lifetime). Since retinoblastoma is a childhood disease, there is no need to construct an age distribution function to calculate the *N_P_* parameter. We can find the *N_P_* parameter by summing all cases of the disease in the considered group of patients. Recall that some cases of cancer in older people are not recorded in the incidence statistics (due to the death of patients from old age), and, in this case, the *N_P_* parameter can only be computed using the age distribution function.

For a model with a sequence of key events (for composite age distribution), the approximation allows calculating the optimal parameter *n_g_* (the number of cells in the group affected by the total carcinogenic event). For the female form of retinoblastoma, *n_g_* = 750,000. For the male form of retinoblastoma, *n_g_* = 130,000. Thus, from the point of view of this model, one oncogenic event affects not a single cell, but several hundred cells.

The maxima of all considered theoretical uni-modal age distributions lie in the age range from 1 to 2 years (here we do not take into account models with a bimodal distribution, for which the distribution function has two local maxima, widely spaced along the time axis).

Not all considered models demonstrate, when optimized, the presence of a clearly expressed minimum of the age distribution function for *T_S_* parameter (latent period of the disease). In four cases, the minimum of the error function is reached when *T_S_* is negative. A negative *T_S_* means we can diagnose cancer before the key event that triggers the disease has occurred. Therefore, in such cases, we enter the value *T_S_* = 0 into the results table.

A negative time *T_S_* appears for two key events (*z* = 2) in a complex mutation model and in model with a sequence of key events (for female and male retinoblastoma). This, in particular, tells us that there is no need to consider models in which the number of key events is more than two – for them the time *T_S_* will also be negative.

The best approximation accuracy (for male and female forms of retinoblastoma) is shown by a model with a sequence of key events (see Table 6.1, lines with *z* = 1.56). Unlike mutational models, this model has four free parameters for optimization (here we also optimize *n_g_* parameter). By using more free parameters, we can more accurately fit the approximation function to real data.

It is noteworthy that this model appears to be the best for both female and male forms of retinoblastoma (the second place in the accuracy of approximation for female and male retinoblastoma is taken by different models).

This model shows the following parameters *T_S_* (theoretical latent period of the disease) *T_S_* = 3.6 months for female retinoblastoma; *T_S_* = 8.5 months for male retinoblastoma. The TS parameter gives us the minimum possible age at diagnosis. The minimum possible age at diagnosis is TS minus the duration of pregnancy (8.72 months). Thus, for male and female retinoblastoma, the model assumes that the disease can be detected at birth – for both hereditary and non-hereditary retinoblastoma.

The mean age at diagnosis for this model is: 14.8 months (hereditary retinoblastoma) and 33.6 months (non-hereditary retinoblastoma) for female retinoblastoma. For male retinoblastoma, the theoretical mean age at diagnosis is 16.ø months months (hereditary retinoblastoma) and 34.7 months (non-hereditary retinoblastoma). We consider the age from the moment of fertilization of the mother’s egg (this is age from birth plus 8.72 months of pregnancy).

The second place in terms of approximation accuracy is taken by the model of one carcinogenic event with two different latencies for female form of retinoblastoma and the complex mutational model for male form of retinoblastoma.

The first of these models (the female form of retinoblastoma), according to the results of approximation, gives very close theoretical values of the mean age at diagnosis for hereditary and non-hereditary forms of the disease (25.9 and 26.5 months respectively). In practice, hereditary retinoblastoma is found at an earlier age.

It should be understood here that when calculating the theoretical mean age of diagnosis, we assume that the frequency of medical examinations of patients is approximately the same for any age. In reality, of course, this is not the case. Both doctors and parents today know that if the parents have retinoblastoma, the child will get the disease with about 50% probability. That is, the risk of illness in a child is very high. Therefore, for hereditary cases, ophthalmologic examinations begin at a very early age and are carried out more frequently. Therefore, the mean age at diagnosis should be biased towards the time of birth (relative to the theoretical value).

As drawbacks of the second of the above models (complex mutational model with compound age distribution (male form of retinoblastoma), we can note the following facts.

First, this model assumes that key events occur in the cell, and not with one of the two alleles of Rb gene (which are in the cell). For the case of two alleles, we built a special model of retinoblastoma – the third mutational model, which (contrary to expectations) shows worse results compared to a complex mutational model (for both female and male forms of retinoblastoma). This fact may tell us that the key events leading to cancer are not mutations of Rb gene allele.

Secondly, for the male form of retinoblastoma, a complex mutational model gives a latency period, which is *T_S_* = 12.2 months. This means that the earliest age we can diagnose retinoblastoma is 3.48 months – before this age (according to the obtained model data) there can be no cases of retinoblastoma (we count the patient’s age from the time of fertilization of the mother’s egg, so the time of birth is *t*_0_ = 8.72 months; the minimum possible age for registration of retinoblastma is 12.2 – 8.72 = 3.48 months). This is contrary to practical observation – cases of retinoblastoma are also recorded in one-month-old children.

Based on the above, from all considered theoretical models of retinoblastoma, we must choose a model with a sequence of key events, which gives the best accuracy of approximation of real data. But it is also necessary to additionally check all the described models on other sets of real data on retinoblastoma available in the databases of large international cancer registries (European, English, Chinese and other cancer registries).

Also, when modeling age distribution, it is desirable to consider separately hereditary and non-hereditary forms of retinoblastoma. This will allow us to more accurately match theoretical models with real data.

It would be useful and interesting if scientists from other countries with access to national cancer statistics would carry out theoretical studies of this kind.

Saint Petersburg

2021

## References

[1] Knudson A.G. Mutation and Cancer: Statistical Study of Retinoblastoma. Proc. Nat. Acad. Sci. USA. Vol. 68, No. 4, April 1971. https://www.pnas.org/content/68/4/820

[2] Dryja T.P., Friend S., Weinberg R.A. Genetic sequences that predispose to retinoblastoma and osteosarcoma. Symp. Fundam. Cancer Res., 39, 1986.

[3] Rushlow D.E., Mol B.M., Kennett J.Y., Yee S., Pajovic S., Th?riault B.L., et al. Characterisation of retinoblastomas without RB1 mutations: Genomic, gene expression, and clinical studies. The Lancet Oncology. 14, 2013.

[4] Paquette L.B., Jackson H.A., Tavare C.J., Miller D.A., Panigrahy A. In Utero Eye Development Documented by Fetal MR Imaging. American Journal of Neuroradiology, 30(9), Oct. 2009. https://doi.org/10.3174/ajnr.A1664

[5] Groot A.L., Lissenberg-Witte B.I., Rijn L.J., Hartong D.T. Meta-analysis of ocular axial length in newborns and infants up to 3 years of age. Survey of Ophthalmology, June 28, 2021. https://doi.org/10.1016/j.survophthal.2021.05.010

[6] Fledelius H.C., Christensen A.S., Fledelius C. Juvenile eye growth, when completed? An evaluation based on IOL-Master axial length data, cross-sectional and longitudinal. Acta Ophthalmologica, Vol. 92, Issue 3, May 2014. https://doi.org/10.1111/aos.12107

[7] Bremner R., Sage J., The origin of human retinoblastoma. Nature, Vol. 514, Issue 3, September 2014.

[8] Curcio C.A., Sloan K.R., Kalina R.E., Hendrickson A.E. Human Photoreceptor Topography. The Journal of Comparative Neurology, February 1990. https://doi.org/10.1002/cne.902920402

[9] Tetearing A.N. Cancer models and statistical analysis of age distribution of cancers. 2021. https://dx.doi.org/10.2139/ssrn.3841599

[10] Tetearing A.N. Theory of populations. SSO Foundation, Moscow, 2012.

[11] National Cancer Institute and US Department of Health and Human Services, SEER As a Research Resource. National Institute of Health, 2010.

[12] Tetearing A.N. Approximation of the age distribution of cancer incidence using a mutational model. 2021. https://doi.org/10.1101/2021.07.06.451349

[13] Tetearing A.N. Using the solution of the Bertalanffy equation as a function of tissue growth for the approximation of age distribution of urinary bladder cancers. 2021.

[14] United States Census Bureau https://www.census.gov/

[15] National Cancer Institute, Surveillance Epidemiology and End Results Overview of the SEER Program. https://seer.cancer.gov/about/overview.html

[16] Draper G.J., Sanders B.M., Brownbill P.A., Hawkins M.M. Patterns of risk of hereditary retinoblastoma and applications to genetic counselling. Br. J. Cancer, 66, 1992. https://doi.org/10.1038/bjc.1992.244

